# A genetically defined injured C-fibre nociceptor associated with neuromas drives chronic spontaneous neuropathic pain

**DOI:** 10.1101/2025.09.30.679489

**Authors:** Xiangsunze Zeng, Rosa I. Martinez-Garcia, Shamsuddin A. Bhuiyan, Keunjung Heo, Natalie MacKinnon-Booth, Elke Bentley, Emily Shea, Kangling Wu, Omer Barkai, Bruna Lenfers Turnes, Selwyn S. Jayakar, Barbara Gomez-Eslava, Mustafa Q. Hameed, Celine Santiago, Karina Lezgiyeva, Emmanuella Osei-Asante, Ivan Furfaro, Alexander Rotenberg, Benjamin R. Johnston, Brian J. Wainger, William Renthal, Stéphanie P. Lacour, David D. Ginty, Clifford J. Woolf

## Abstract

Spontaneous pain is a common but poorly understood consequence of peripheral nerve injury^1–3^, including injuries that lead to the formation of neuromas^4,5^. We developed a chronic neuroma model for measuring spontaneous pain-related behaviours in mice, which revealed that limb flicks - emerging predominantly 2 months post-injury - reflect spontaneous paroxysmal pain. Ectopic activity of injured dorsal root ganglia (DRG) sensory neurons whose axonal endings terminate within the neuroma drives this spontaneous pain. *In vivo* imaging showed that a subset of small-diameter DRG sensory neurons are the source of spontaneous neural signals emanating from the neuroma, and these spontaneously active neurons are distinct from the intact larger diameter sensory neurons that mediate stimulus-evoked mechanical allodynia from spared nerves. Cell-type–specific gain- and loss-of-function studies identified a genetically- and functionally-defined subtype of small-diameter C-fibre nociceptors whose injured axons in neuromas drive spontaneous limb flicks/neuropathic pain. These findings establish the neurobiological basis of spontaneous pain enabling targeted pain management strategies and define a cellular and mechanistic separation between spontaneous and evoked neuropathic pain.

## Introduction

Peripheral nerve injury is a major cause of chronic neuropathic pain, resulting in persistent suffering and disability. Spontaneous pain is among the most common and debilitating clinical feature of neuropathic pain^1–3^, yet its underlying mechanisms remain largely unknown. Most preclinical research emphasizes stimulus-evoked withdrawal responses^6^, which poorly captures the chronic and fluctuating nature of spontaneous neuropathic pain. This knowledge gap stems from the fact that spontaneous pain arises sporadically and unpredictably, posing challenges for comprehensive quantitative investigations^7^ and mechanistic discovery.

Existing measures of spontaneous pain are inadequate. For example, the grimace scale performs well in acute pain settings but not for neuropathic pain^8^. Conditioned place preference can indicate ongoing pain^9^, but fails to capture the paroxysmal nature of spontaneous pain—characterized by sudden, episodic bouts—which is a hallmark in patients with peripheral nerve trauma who develop neuromas^10^. Consequently, studies have relied on intermittent pain-like behaviours such as paw biting or guarding to infer spontaneous paroxysmal pain after nerve injury^11,12^. These measures, however, typically depend on short-term video observations and do not provide a comprehensive view of sporadic and unpredictable pain episodes over prolonged periods.

Neuropathic pain research has predominantly centered on acute and subacute post-nerve injury (e.g., 1–4 weeks), leaving its chronic features relatively underexplored^13^, even though chronic pain is the major clinical issue. Furthermore, the cellular sources of aberrant activity are unclear: early electrophysiological studies demonstrated that both injured and neighboring uninjured sensory fibres generate ectopic discharges soon after axotomy^12,14,15^, but the genetically defined and physiologically distinct sensory subtypes that drive chronic spontaneous neuropathic pain have not been identified. Finally, a longstanding clinical puzzle is why patients with comparable peripheral nerve insults experience different outcomes, with some developing severe chronic pain while others exhibiting moderate or no pain^4,5,16^. Although risk factors, including genetic predisposition, psychosocial context, prior injury, age, and sex, have been proposed^16,17^, the cellular and molecular bases for this variability are still elusive.

By integrating long-term home-cage behavioural studies with a novel measure of spontaneous pain, *in vivo* calcium imaging, cell-type–specific optogenetic and chemogenetic manipulations, and transcriptomic profiling in mice, we now identify injured axons of a genetically and functionally defined C-fibre nociceptor subtype as the driver of chronic spontaneous neuropathic pain in animals with painful neuromas. Our findings also reveal a clear cellular and mechanistic distinction between spontaneous and evoked neuropathic pain.

## Results

### Painful neuromas drive chronic spontaneous pain after peripheral nerve injury

We sought to understand the mechanistic basis of spontaneous neuropathic pain by analysing defined spontaneous behaviours in mice using a machine-learning–based platform^18^ following different types of sciatic nerve injury (Extended Data Fig. 1a,b). Guarding, biting/licking, scratching, rearing, and grooming behaviours were monitored using 30-minute recording sessions conducted once per week for up to two months after injury. All of these behaviours remained unchanged post-injury, except for guarding, where mice lift and hold the paw ipsilateral to the nerve injury, which increased relative to sham controls from one week after the injury (Extended Data Fig. 1c), in line with previous studies^11,12^.

While the conventional behavioural measures did not provide a reliable measurement of spontaneous pain, the long-term videography analysis revealed an unexpected “limb flick” behaviour that was very rare in the first 4 weeks after the nerve injury, but then progressively increased, becoming most prominent at 8 weeks post-injury across all the nerve transection injury models (Extended Data Fig. 1d). This spontaneous limb flick behaviour was characterized as having a stereotyped pattern of rapid, repetitive hindlimb shaking (Supplementary Video 1). The incidence of limb flicks scaled with injury severity, being most pronounced after a full sciatic nerve transection (SNT) and intermediate in the spared nerve injury (SNI) model, where the common peroneal and tibial nerves are injured and the sural nerve remains intact (Extended Data Fig. 1a,d). Limb flicks were also observed after a sural nerve transection (SuNT) (Extended Data Fig. 1a,d), which selectively damages sensory fibres^19^, and they were suppressed by gabapentin (Extended Data Fig. 2a), supporting that this behaviour likely reflects pain and not motor dysfunction. By contrast, limb flicks were absent in acute nociceptive or inflammatory pain models (Extended Data Fig. 2b). These observations suggest that spontaneous limb flicks arise from chronic pathological changes in injured sensory nerve fibres.

Because the limb flick is a chronic pain-associated behaviour that emerged most robustly in the SNT model, we focused on this nerve injury model for subsequent analyses. To move beyond short-term measures of limb flicks, we established a quantitative framework for long-term home-cage monitoring. Given the stereotyped high-frequency muscle activity underlying limb flicks, we implemented a wireless electromyography (EMG) system using electrodes placed in the vastus lateralis, the largest quadriceps muscle, coupled with video recording, to reliably capture injured hindlimb behaviours over extended time periods (Fig. 1a). The EMG system faithfully detected repetitive limb flick oscillations (Supplementary Video 1), with an average peak frequency of ∼13 Hz and most events lasting <1 second (Extended Data Fig. 3a–c), without any apparent motor impairment from the recording electrode implantation (Extended Data Fig. 3d). To determine whether prolonged monitoring reduces behaviour variability, we performed continuous 24-hour recording at 8 weeks post-SNT and assessed intra-animal variation by binning the data into time windows of increasing length. As the window size increased, variability progressively declined, and extending the recording period up to 48 hours did not yield a further reduction when compared with 24 hours (Fig. 1b). These results demonstrate that 24-hour monitoring provides a stable measure of limb flicks and also reveal a circadian component to this spontaneous neuropathic pain behaviour^20^.

**Fig. 1.**
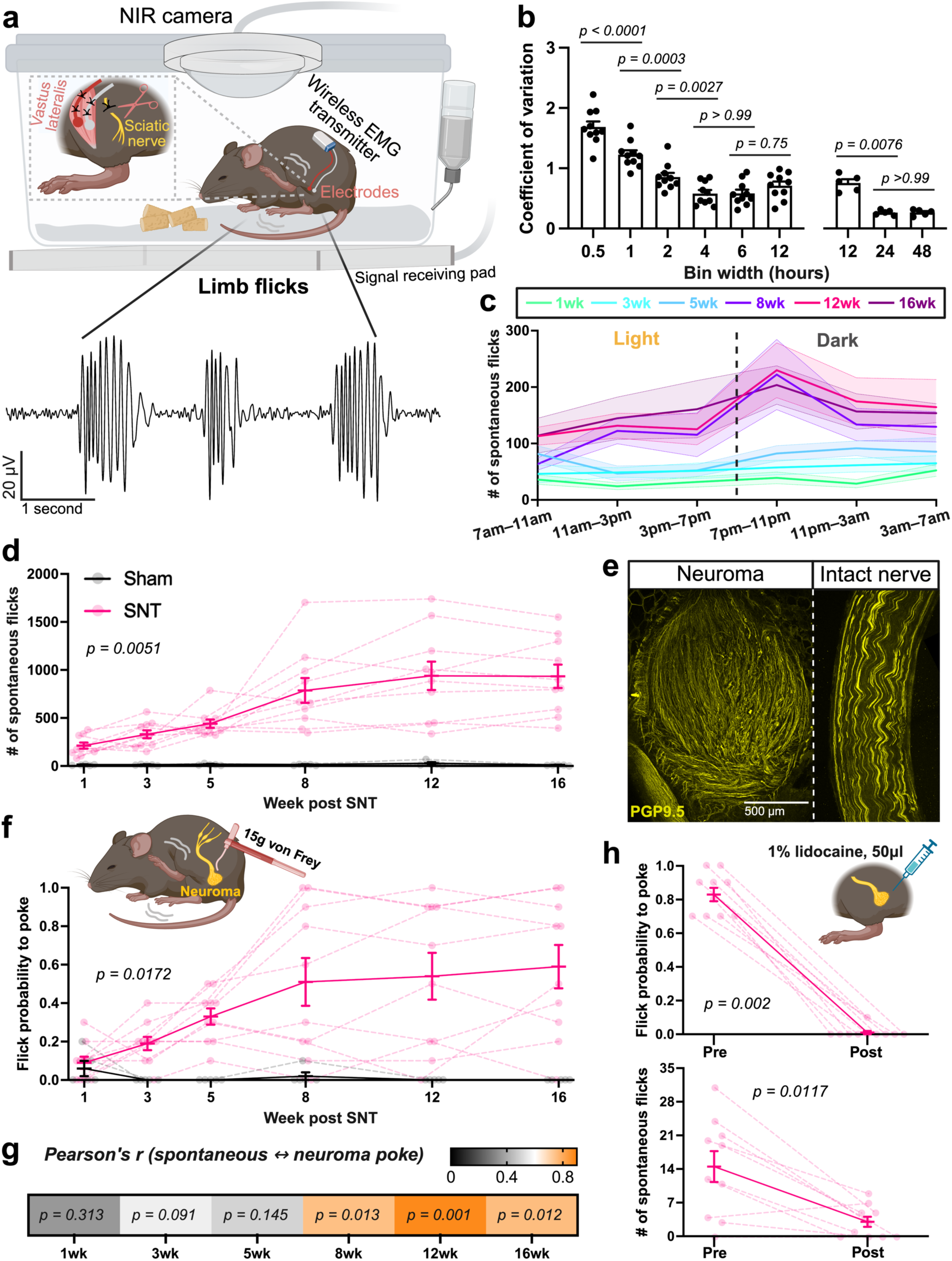
| Neuromas drive chronic spontaneous neuropathic pain. **a**, Schematic of the home-cage EMG/video recording system (top) and representative EMG trace of limb flick bouts (bottom) after SNT. **b**, Coefficient of variation (CV) of spontaneous limb flick counts following binning of 24-hour recordings into different time windows (0.5, 1, 2, 4, 6, and 12 hours). Each dot represents one recording from an SNT mouse (n = 10). A subset of five mice was recorded for 6 consecutive days to compute CV across extended 12-, 24-, and 48-hour windows. Data are mean ± SEM. *P* values from one-way ANOVA with Tukey’s multiple comparisons within each dataset. **c**, Spontaneous limb flick counts quantified in 4-hour bins over 24 hours at indicated time points post-SNT. Light phase: 7:00 a.m.–7:00 p.m.; dark phase: 7:00 p.m.–7:00 a.m. Solid lines and shaded bands represent mean ± SEM across n = 10 SNT mice. **d**, Total 24-hour spontaneous flick counts in individual mice after sham surgery (n = 5) or SNT (n = 10). Data are mean ± SEM; dashed lines connect individual mice. *P* values from two-way ANOVA with Tukey’s multiple comparisons (figure shows the interaction effect of sham vs SNT over time). **e**, Confocal images of a neuroma (8 weeks post-SNT) and contralateral intact nerve labelled with a neuronal axonal marker PGP9.5. **f**, Flick probability in response to transcutaneous neuroma poke after sham surgery or SNT in the same mice shown in (**d**). **g**, Pearson’s correlation between spontaneous flick counts (**d**) and flick probability (**f**) at indicated time points post-SNT. **h**, Flick probability to poke (top) and spontaneous flick counts over 30 minutes (bottom) before and after local lidocaine injection at the neuroma site (8–12 weeks post-SNT, n = 10 high-pain mice). Data are mean ± SEM; dashed lines connect individual mice. *P* values from two-sided Wilcoxon matched-pairs signed-rank test.

Using 24-hour recordings, we tracked limb flicks longitudinally. Group averages rose gradually, reaching a peak around 8 weeks post-SNT and remaining at a plateau thereafter, with most increases occurring during the dark phase (7 p.m.–7 a.m.) (Fig. 1c,d). Between mice, the flick prevalence varied, with some mice displaying dramatic increases and others remaining at moderate or low levels throughout the 16-week post-SNT period (Fig. 1d). Unlike the intra-animal variability attributable to recording length, this inter-animal variability likely reflects biological differences and is reminiscent of the different pain phenotypes in patients who develop painful or non-painful neuromas after peripheral nerve injury or an amputation^4,5^. Histology confirmed the formation of terminal neuromas proximal to the nerve transection, appearing as bulbous structures filled with exuberant but disorganized axons within fibrotic tissue by 8 weeks post-SNT (Fig. 1e), as described previously^21^. To test whether the neuromas trigger the limb flicks, we used a transcutaneous poke test, analogous to the clinical Tinel sign evoked by gentle tapping the skin over a peripheral nerve^22^. Our poke test involved indenting the skin overlying the SNT lesion with a 15 g von Frey filament to mechanically stimulate the underlying neuroma (hereafter referred to as “transcutaneous neuroma poke”). This stimulus increasingly induced limb flicks over time across the nerve injury models, with the strongest responses in SNT animals, and minimal responses in sham controls or when applied to the side contralateral to the SNT surgery (Extended Data Fig. 4a and Supplementary Video 2). EMG implantation did not affect this outcome (Extended Data Fig. 3e). We also assessed flick probability to the transcutaneous neuroma poke in the same mice that were used for the 24-hour spontaneous flick recordings. This analysis revealed comparable inter-animal variability as spontaneous limb flicks (Fig. 1f) and also demonstrated a strong positive correlation in the chronic phase: mice with more spontaneous limb flicks were also highly sensitive to the poke test, whereas those with few spontaneous limb flicks showed little or no flick response to a poke (Fig. 1g). To test causality, we injected lidocaine locally at the neuroma site, a standard clinical intervention^23^. This virtually abolished the poke-induced flicks in mice with a high neuroma pain behaviour. In addition, lidocaine treatment markedly reduced spontaneous flicks (Fig. 1h), indicating that activity originating in neuromas is the major driver of this chronic spontaneous pain behaviour.

To quantify the incidence of the high-pain phenotype, we performed the transcutaneous neuroma poke test in 144 mice at 8–12 weeks post-SNT. Thirty-two percent of SNT mice exhibited a high response probability (≥0.7; hereafter referred to as high-pain), twenty-seven percent exhibited low probability (≤0.3; low-pain), and forty-one percent displayed an intermediate pain phenotype (0.4–0.6; medium-pain), with no sex differences or correlation with the neuroma size (Extended Data Fig. 4b,c). Finally, neither poke-induced nor spontaneous limb flicks required the mechanosensitive channel Piezo2, as evaluated in Piezo2 conditional knockout mice (Extended Data Fig. 5).

### Injured small-diameter sensory axons exhibit ectopic activity arising from their terminals in the neuroma

To investigate the neuronal basis of the spontaneous pain, we performed *in vivo* calcium imaging from the L4 dorsal root ganglia (DRG) 8–12 weeks after SNT using *Vglut2^Cre^::GCaMP6f* mice, in which primary DRG sensory neurons across all subtypes are labelled. The neuroma or contralateral intact nerve was surgically exposed to allow direct stimulation with von Frey filaments of varying force (Fig. 2a). Poking neuromas induced calcium responses almost exclusively in small-diameter neurons (<25 µm soma diameter), whereas stimulation of the contralateral intact nerve rarely activated neurons of any size, with the 1.4 g filament producing the most discriminating response (largest effect size determined by Cohen’s d) (Fig. 2b,c). Neuroma-poke responses also varied across animals: the response amplitude of small-diameter neurons induced by the 1.4 g probe stimulus differed substantially across mice and was positively correlated with their behavioural pain responsiveness, assessed by transcutaneous neuroma poke immediately prior to the imaging (Fig. 2d), supporting a peripheral contribution to the differential pain outcomes in individual mice.

**Fig. 2.**
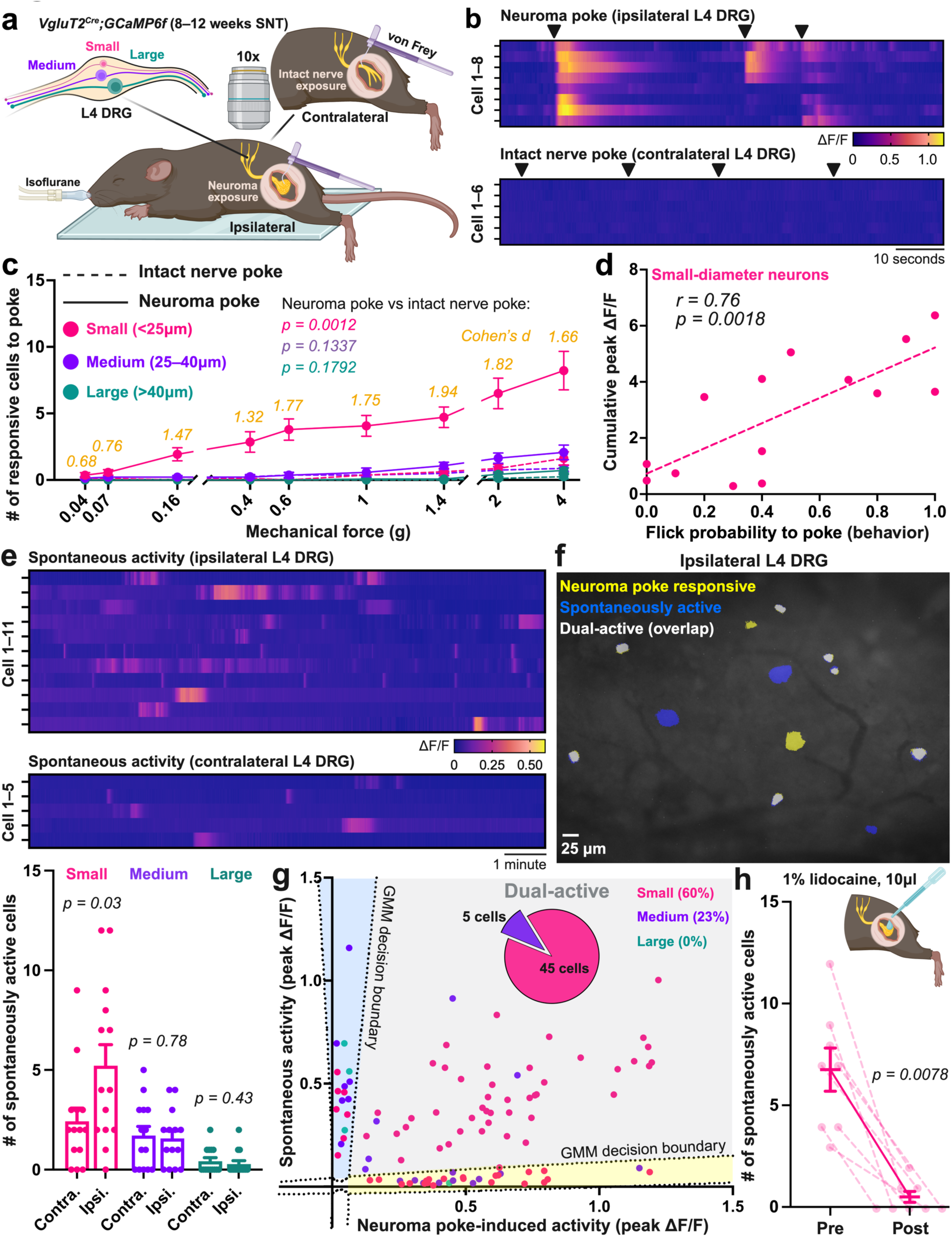
| Small-diameter DRG neurons respond to neuroma poke and exhibit ectopic spontaneous activity. **a**, Schematic of the *in vivo* DRG imaging setup. **b**, Calcium activity heatmaps from individual DRG neurons from one representative mouse in response to a 1.4 g von Frey filament applied to the neuroma or to the contralateral intact nerve. Arrowheads indicate stimulus timing. For intact nerve poke, cells were defined by responses to plantar paw stimulation with the same filament, as no cells showed detectable responses to the nerve poke. **c**, Quantification of responsive cells by soma diameter, small (<25 μm), medium (25–40 μm), and large (>40 μm), across a range of mechanical forces applied with von Frey filaments. Data are mean ± SEM from 14 ipsilateral and contralateral L4 DRG (n = 14 mice). *P* values from two-way ANOVA with Tukey’s multiple comparisons (figure shows the interaction effect of neuroma poke vs intact nerve poke across mechanical forces). Cohen’s d effect sizes are shown for small-diameter neurons (neuroma poke versus intact nerve poke). **d**, Pearson’s correlation between the cumulative peak ΔF/F of responsive small-diameter DRG neurons to neuroma poke (1.4 g von Frey) and behavioural flick probability to transcutaneous neuroma poke. Each dot represents one mouse. **e**, Representative heatmaps of spontaneous calcium activity from individual neurons from one representative mouse in ipsilateral and contralateral L4 DRG (top), and quantification of spontaneously active cells of each size during 30-minute recordings (bottom). Data are mean ± SEM; each dot represents one L4 DRG (n = 14 DRG from 14 mice). *P* values from two-sided paired t-tests. **f**, Example overlay of cells spontaneously active (blue) and cells responsive to neuroma poke (yellow); dual-active cells are overlaid (white). **g**, Classification of cells solely responsive to neuroma poke, solely spontaneously active, or dual-active, based on peak activity amplitude determined by Gaussian Mixture Model (GMM) boundaries (dashed lines). Percentages indicate the proportion of dual-active cells relative to the total number of cells of each size. Pie chart depicts the cell distribution within the dual-active category. Each dot represents one cell. **h**, Number of spontaneously active small-diameter cells before and 5 minutes after topical application of lidocaine to the neuroma. Data are mean ± SEM; dashed lines connect ipsilateral L4 DRG from individual mice (n = 8). *P* value from two-sided Wilcoxon matched-pairs signed-rank test.

We next assessed spontaneous activity in the same DRG and found that the L4 DRG ipsilateral to the neuroma contained more spontaneously active cells than the contralateral L4 DRG containing neurons with an intact sciatic nerve, specifically within the small-diameter population (Fig. 2e). Comparing the ipsilateral spontaneously active neurons with those responsive to a neuroma poke showed that the majority—both spontaneously active and those responsive to a poke—were small-diameter neurons (Fig. 2f,g). Moreover, topical lidocaine applied directly at the neuroma largely abolished the spontaneous activity (Fig. 2h), demonstrating that the origin of this spontaneous neural activity is in the neuroma rather than the DRG. These findings establish injured small-diameter sensory axons as the principal source of ectopic activity and the likely drivers of spontaneous neuropathic pain.

### Pain arising from injured versus uninjured fibres requires distinct sensory neuron cell types

Peripheral nerve injury induces ectopic activity in both injured and adjacent uninjured fibres across different sensory neuron subtypes^12,14,24,25^. We next asked whether activity originating from injured fibres in neuromas and uninjured fibres jointly contribute to a common pain phenotype or act through distinct mechanisms. Because the entire sciatic nerve is severed in the SNT model, we assessed mechanosensitivity of the medial plantar paw which is innervated by the intact saphenous nerve ^26^. Tactile allodynia associated with the medial plantar paw emerged 4 weeks post-SNT (Fig. 3a), well before spontaneous limb flicks became prevalent. No correlation was found between the intact saphenous nerve-evoked pain and neuroma pain intensity (Fig. 3b), pointing to independent mechanisms. Direct imaging of DRG activity confirmed this separation: completely distinct populations of sensory neurons were engaged in the two pain models. Medium- and large-diameter DRG neurons were the primary responders to medial plantar stimulation, whereas small-diameter neurons were predominately associated with neuroma stimulation (Fig. 3c–f; and Supplementary Video 3). Locally silencing the neuroma or saphenous nerve endings with lidocaine selectively inhibited responses initiated from each region, respectively (Fig. 3g,h), providing evidence that pain arising from injured and uninjured fibres is initiated through distinct mechanisms.

**Fig. 3.**
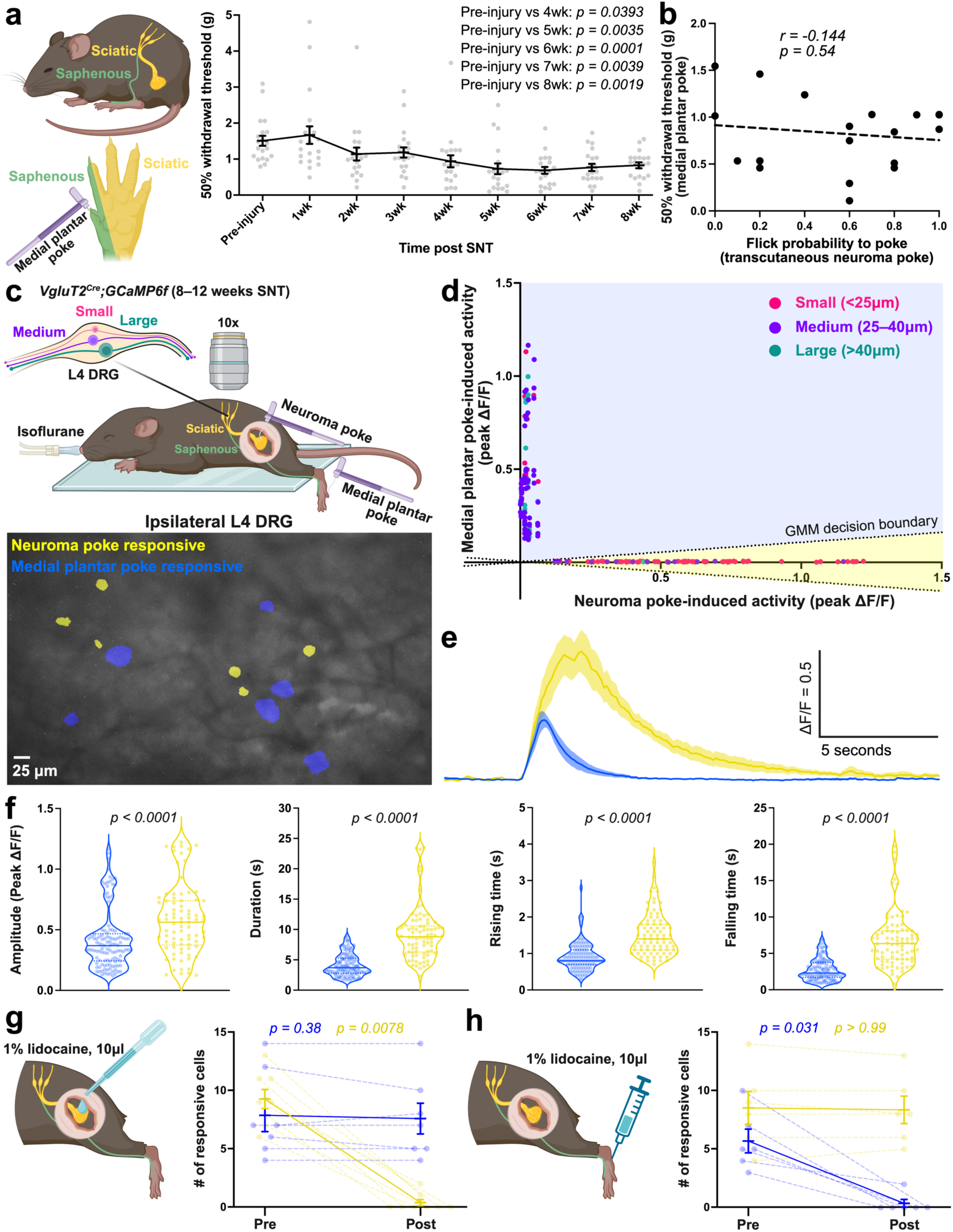
| Distinct DRG neuron populations mediate pain from intact versus injured sensory fibres. **a**, Schematic of the plantar von Frey assay targeting the medial plantar area innervated by the intact saphenous nerve (left) and quantification of the 50% withdrawal threshold (right). Each dot represents one mouse (n = 20). Data are mean ± SEM. *P* values from one-way ANOVA with Tukey’s multiple comparisons. **b**, Pearson’s correlation between the 50% withdrawal threshold from medial plantar poke and flick probability to transcutaneous neuroma poke at 8 weeks post-SNT. Each dot represents one mouse. **c**, Schematic of the *in vivo* L4 DRG calcium imaging setup for neuroma or medial plantar poke with a 1.4 g von Frey filament (top), and an example overlay showing cells responsive to neuroma poke and medial plantar poke (bottom). **d**, Classification of cells responsive to neuroma poke or medial plantar poke based on peak ΔF/F amplitude determined by GMM boundaries (dashed lines). Each dot represents one cell; no dual-responsive cells were observed. **e**, Calcium traces from neuroma poke-responsive (yellow) and medial plantar poke-responsive (blue) cells. Shaded bands indicate mean ± SEM from n = 5 mice (neuroma poke) and n = 4 mice (medial plantar poke). **f**, Quantification of response amplitude, duration, rising time, and falling time for each group. Each dot represents one cell. *P* values from two-sided Mann–Whitney tests. **g**,**h**, Number of cells responsive to a neuroma poke (yellow) or a medial plantar poke (blue) before and after topical lidocaine application to the neuroma (**g**) or intraplantar injection (**h**). Tests were performed 5 minutes after lidocaine administration. Data are mean ± SEM; dashed lines connect individual mice (n = 8 for neuroma application, n = 6 for medial plantar injection). *P* values from two-sided Wilcoxon matched-pairs signed-rank tests.

### A genetically defined subtype of C-nociceptor axons in neuromas drives limb flicks

Transcriptomic studies have identified at least 16 genetically distinct DRG sensory subtypes^27–31^ with unique physiology and morphology^28^. We performed single-nucleus RNA sequencing (snRNA-seq) of DRG from mice 10 weeks post-SNT and confirmed the presence of all these cell types in both high- and low-pain conditions (Fig. 4a and Extended Data Fig. 6a). To identify specific sensory subtypes whose activation may drive spontaneous limb flicks, we conducted an optogenetic gain-of-function screen, using transgenic mouse lines that collectively encompass the majority of sensory subtypes with soma diameters <25 µm ^28^. Because neuromas reside deep within fibrotic tissue, we adapted a miniaturized LED cuff ^32^ for localized epineural activation of sensory axons, either in neuromas or contralateral intact nerves (Fig. 4b and Extended Data Fig. 7). In mice expressing opsins in a subset of small-diameter C-fibres marked by *Sstr2* expression (SSTR2; *Sstr2^T2a-CreER^;Calca-FlpE;R26^LSL-FSF-ReaChR::mCitrine^*), a single 10-ms 470-nm pulse delivered to the neuroma induced limb flicks and, to a lesser extent, other pain-related behaviours (vocalization, jump, licking, and avoidance, either individually or in combination)^7^, with significantly lower activation thresholds than those needed to evoke behavioural responses on the contralateral intact nerve (Fig. 4c,d). The nomenclature for this cell type varies across the literature, depending on the response modality assessed in intact tissue: C-Heat thermonociceptor (C-Heat [SSTR2]) from skin^28^; C-high threshold mechanoreceptors (C-HTMR) from colon^33^, lineage subtype (CGRP-α)^29^, marker gene expression in mice (*Calca* + *Sstr2*)^30^, hierarchical transcriptomic clustering within the peptidergic branch (PEP1.1) ^31^, and putative human homologues (C-PEP.TAC1/CACNG5)^34^. Here, we refer to this genetically defined, neuroma-hypersensitive C-fibre subtype as ‘Neuroma C-Nociceptors (SSTR2)’. In contrast, Aδ-HTMR/Heat (SMR2) neurons, which are myelinated mechanonociceptors^35^, elicited behavioural responses when intact nerves were stimulated but rarely when neuromas were activated (Fig. 4c,d). This discrepancy likely reflects the preferential loss of the axons of these Aδ-HTMR/Heat (SMR2) neurons in neuromas (Extended Data Fig. 8). Other subtypes showed little or no nocifensive behavioural responses when activated: activating Aδ-low-threshold mechanoreceptors (Aδ-LTMRs), C-LTMRs, and C-HTMR/Heat (CYSLTR2/SST) neurons elicited no consistent behaviours from either neuromas or intact nerves, whereas stimulating C-HTMR/Heat (MRGPRA3), C-HTMR/Heat (MRGPRB4), C-HTMR/Heat (MRGPRD), C-Cold (TRPM8) axons with strong optical power occasionally evoked avoidance, guarding, or licking behaviours from either site, but rarely induced limb flicks (Fig. 4c,d; and Supplementary Video 4).

**Fig. 4.**
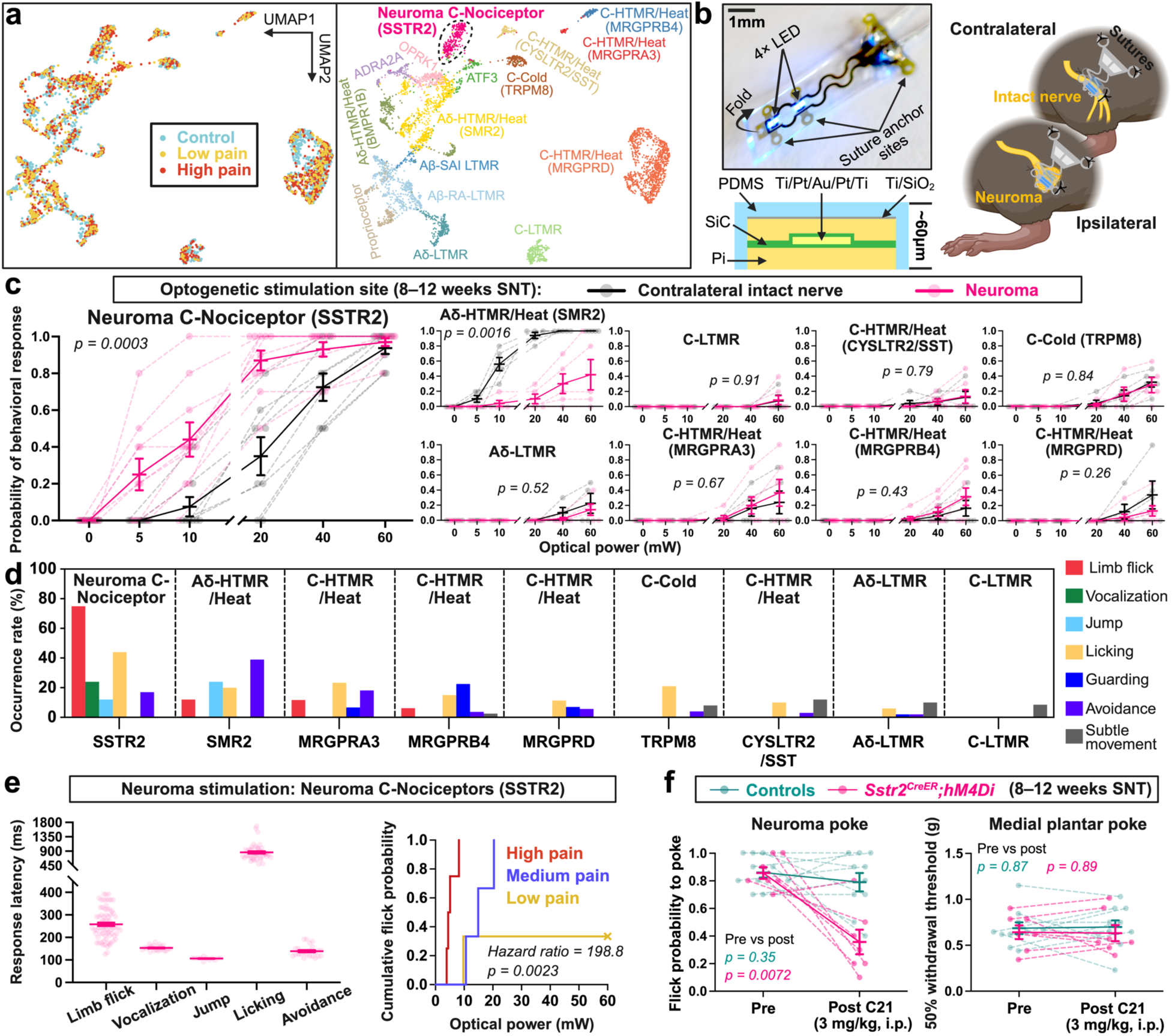
| Neuroma C-Nociceptors (SSTR2) drive limb flicks. **a**, UMAP projections of DRG snRNA-seq showing nuclei from contralateral DRG (control), ipsilateral DRG from low-pain, and high-pain mice (left) and annotated sensory-neuron subtype identities (right). 10 weeks post-SNT mice were used for snRNA-seq. **b**, Photograph of the miniaturized blue LED cuff (top), schematic cross-section of the interconnects showing layered materials (bottom left; softness and flexibility were achieved by miniaturizing metal interconnects and incorporating soft polymer layers; see Extended Data Fig. 7 for details), and illustration of the implantation site for neuroma or intact sciatic nerve stimulation (right). **c**, Optogenetic activation of genetically defined neuronal subtypes (see Methods for the transgenic lines targeting each subtype). Optical power versus response probability for individual transgenic lines (10 trials per animal; single 10 ms pulse). Data are mean ± SEM; dashed lines connect individual mice. *P* values from two-way ANOVA with Tukey’s multiple comparisons (figure shows the interaction effect of neuroma versus intact nerve stimulation across optical powers). **d**, Histograms showing the occurrence of different behaviours evoked at maximum optical neuroma stimulation (60 mW) using single 10-ms pulses. Note that multiple behaviours could occur in a single trial. **e**, Left: response latencies of individual behaviours relative to stimulus onset; data are mean ± SEM, each dot represents a single trial across n = 10 mice. Right: Cumulative flick probability (1 − Kaplan-Meier estimate) as a function of optical power for high-, medium-, and low-pain mice determined by transcutaneous neuroma poke. Cross mark at 60 mW indicates right-censoring for animals who did not show a flick at the maximum optical power. **f**, Limb flick probability to transcutaneous neuroma poke (left) and 50% withdrawal threshold from medial plantar poke before and after intraperitoneal (i.p.) injection of compound 21 (C21) in *Sstr2^CreER^;hM4Di* mice (n = 7) for selective chemogenetic silencing of Neuroma C-Nociceptors (SSTR2) and littermate controls (*Sstr2^CreER^* alone or *hM4Di* alone, n = 10). Data are mean ± SEM; dashed lines connect individual mice. *P* values from two-sided paired t-tests.

Anatomical analyses revealed denser Neuroma C-Nociceptor (SSTR2) fibres in neuromas than in contralateral intact nerves, but a reduced presence of Aδ-HTMR/Heat (SMR2), C-HTMR/Heat (MRGPRA3), and C-HTMR/Heat (MRGPRD) nerve fibres, with C-HTMR/Heat (MRGPRD) axons almost completely absent (Extended Data Fig. 8), likely reflecting cell death in this population after peripheral nerve trauma^36^.

Because activation of the Neuroma C-Nociceptor (SSTR2) injured axons robustly induced limb flicks, we further characterized how their activity relates to different pain-related behaviours and response dynamics. High-frequency or prolonged optogenetic stimulation did not change the incidence of limb flicks or most other behaviours, except audible vocalizing - a response uniquely triggered by activation of the Neuroma C-Nociceptors (SSTR2) - showed an increased occurrence under these stimulation patterns, suggesting a dependence on summation of activity (Extended Data Fig. 9a). Response latency analyses showed that the limb flicks occurred with an average latency of 259 ms, which was longer than the latency to jump or avoid the stimulus but considerably shorter than the latency for self-licking (Fig. 4e, left). These temporal distinctions suggest that limb flicks represent a pain-related behavioural output that is dissociable from rapid withdrawal reflexes^35^ and from delayed coping behaviours. Lastly, we noted from the snRNA-seq analysis that *Scn10a* (encoding voltage-gated sodium channel Na_v_1.8) and *Scn3a* (Na_v_1.3) were selectively upregulated in Neuroma C-Nociceptors (SSTR2) under high-pain conditions (Extended Data Fig. 6b,c), potentially linking these ion channels to hyperexcitability of this population in painful neuromas. Related to this, we analysed the optical stimulation strength required to induce limb flicks in high-, medium-, and low-pain mice. We used a Cox proportional hazards model with optical power as the ordered exposure variable and the first limb flick to optogenetic stimulation as the event. This analysis revealed that the flick probability determined by the transcutaneous neuroma poke test significantly predicted optogenetic threshold (*hazard ratio* = *198.8*, *p* = *0.0023*); high-pain mice flick at a lower optical power, whereas low-pain mice require a higher optical power to elicit flicks (Fig. 4e, right). This result is consistent with increased excitability of Neuroma C-Nociceptors (SSTR2) in high-pain neuromas potentially mediated by upregulation of voltage-gated sodium channels^24^.

We next asked whether activity of the Neuroma C-Nociceptors (SSTR2) is necessary for neuroma-driven pain behaviours. We acutely silenced this neuronal population using chemogenetic inhibition highly specific to the *Sstr2^+^*population (*Sstr2^T2a-CreER^;R26^LSL-hM4Di::mCitrine^*) (Extended Data Fig. 9b). This manipulation significantly suppressed transcutaneous neuroma poke–induced limb flicks in high-pain mice but not in littermate controls (Fig. 4f). Notably, motor function and mechanical allodynia arising from the intact saphenous nerve-innervated area were unaffected (Fig. 4f and Extended Data Fig. 9c), indicating a specific role for activity in injured Neuroma C-Nociceptors (SSTR2) in driving pain arising from the neuroma. These findings define a specific nociceptor subtype whose injured axons become hypersensitive in painful neuromas to drive a chronic spontaneous pain-related behaviour.

## Discussion

The mechanisms of spontaneous neuropathic pain are obscure, largely because of the lack of reliable measures, particularly over chronic timescales. Most mechanistic insights into neuropathic pain have been derived from stimulus-evoked pain arising from uninjured afferents^26^, potentially biasing interpretation toward spared-nerve pathways after injury. By establishing a long-term recording platform, we identified limb flicks as a robust behavioural readout of chronic paroxysmal spontaneous pain and traced their origin to injured axons of a genetically defined sensory subtype within painful neuromas. These findings support the conclusion that the spontaneous and evoked components of neuropathic pain arise from distinct peripheral drivers.

This distinction reframes prevailing models of neuropathic pain and has direct implications for treatment. Prior work indicates that mechanical allodynia mediated by spared nerves after injury is primarily initiated by Aδ/Aβ-LTMRs whose central afferents reroute into nociceptive circuits through dorsal horn plasticity^37,38^, with Piezo2-dependent mechanotransduction playing a central role^39^. By contrast, our model identifies injured Neuroma C-Nociceptors (SSTR2) as the principal drivers of neuropathic pain, and with neuroma mechanical hypersensitivity occurring independently of Piezo2 (Extended Data Fig. 5), which is consistent with minimal expression of Piezo2 in this neuronal subtype^28–31^. Defining the pain transduction mechanisms at neuroma terminals^24^, and how Neuroma C-Nociceptors (SSTR2) engage central circuits to generate pain perception, will advance our understanding of the underlying mechanisms of spontaneous neuropathic pain.

Consistent with the mechanistic separation of spontaneous and evoked pain, both preclinical and clinical evidence suggests that spontaneous pain and stimulus-evoked hypersensitivity are distinct in their pharmacological responsiveness^40–43^. However, current diagnostic frameworks lump these manifestations under a single “neuropathic pain” label, and treatment guidelines recommend single agents without any delineation for evoked or spontaneous pain. This likely contributes to variable therapeutic efficacy through a mechanism–therapy mismatch. Our findings support the need to revisit diagnostic criteria to explicitly distinguish between spared-fibre evoked allodynia and injured-fibre triggered spontaneous pain, and develop specific targeted interventions aligned with the dominant mechanistic driver in individual patients.

A second key implication is that identifying the peripheral driver of spontaneous pain provides a rational framework for cell-targeted intervention^44^. Because Neuroma C-Nociceptors (SSTR2) are both necessary and sufficient to drive neuroma pain, and their selective silencing suppresses neuroma pain without impairing sensory modalities from spared nerves or motor function (Fig. 4f and Extended Data Fig. 9c), therapeutic strategies that act selectively within this population, particularly those enabling localized, subtype-specific delivery^45^, offer a means to alleviate pathological pain while preserving desired sensory function and minimizing central exposure in patients. Although the Neuroma C-Nociceptors (SSTR2) identified here are defined by *Sstr2* expression in mice—a marker with minimal expression in human DRG neurons—the model recapitulates key clinical features of neuropathic pain, suggesting the existence of an evolutionarily conserved homologous human nociceptor subtype, which is supported by transcriptomic observations^34^. Defining the analogous mechanisms operating in humans and identifying molecular access points to suppress neuroma-associated hyperexcitability will be essential for translation.

Beyond symptomatic control, preventing the formation of painful neuromas may be a new treatment strategy. The emergence of high-pain versus low-pain phenotypes that can be decoded at the level of sensory neuron activity (Fig. 2d) points to peripheral heterogeneity in how neuromas are primed after injury. Such variability may reflect differences in stochastic perturbations of gene expression during the early injury response^46,47^, local trophic and inflammatory signaling^48^, and interactions with diverse non-neuronal cell types^48–52^. Defining the determinants that govern painful neuroma maturation during the subacute formation phase may therefore enable early protective strategies that can intercept pathological trajectories before chronic spontaneous pain becomes established, enabling its prevention^44^.

## Methods

### Mice

All experiments were conducted in accordance with the guidelines of the Boston Children’s Hospital Institutional Animal Care and Use Committee (IACUC) under protocols 00001507 and 00002403, or by the Harvard Medical School IACUC under protocol IS00001036-9. Mice were housed under temperature-controlled conditions on a 12-hour light/dark cycle (lights on at 7:00 a.m., off at 7:00 p.m.) with ad libitum access to food and water. Both male and female mice were used in all experiments. The C57BL/6J strain (JAX #000664; The Jackson Laboratory) was used unless otherwise specified, with transgenic strains detailed in the relevant sections below.

### Peripheral nerve injury models

Surgeries were performed on 8–9-week-old mice under aseptic conditions. Anaesthesia was induced and maintained with 2% isoflurane (Patterson Veterinary, #07-890-8115) delivered via a nose cone. Throughout the procedure, body temperature was maintained at 37°C using a heated pad to prevent hypothermia. Sciatic nerve injuries were performed using standardized protocols as follows. 1. The surgical site was shaved, and residual hair was completely removed using Nair. The area was then sterilized with 70% ethanol and Betadine. 2. A small skin incision was made over the lateral thigh, above the biceps femoris. The underlying musculature was carefully separated and retracted to expose the sciatic nerve. 3. For sham controls, the sciatic nerve was gently freed from surrounding connective tissue using curved forceps without causing any damage. For SNT injury, a segment of the sciatic nerve trunk 5 mm above the sural, common peroneal, and tibial nerves was ligated with 6-0 sutures (Ethicon, #K889H) followed by a complete transection proximal to the ligation using microsurgical scissors. An additional 5 mm segment of the distal nerve stump was excised. For SNI injury, the common peroneal and tibial nerve branches were ligated and transected in the same manner, leaving the sural nerve intact. For SuNT injury, the sural nerve branch was ligated and transected in the same manner, leaving the common peroneal and tibial nerve branches intact (Extended Data Fig. 1a). 4. The musculature was then folded back to cover the sciatic nerve with the skin closed with 6-0 sutures. 5. Mice were placed on a 37°C heated pad postoperatively until they fully recovered before being returned to their home cages.

### Automated behavioural screening

The recording setup and analytic platform were identical to those previously described^18^. Briefly, mice were recorded inside a black acrylic enclosure (18 × 18 × 15 cm; length × width × height), which was closed on all sides except for the bottom and positioned on a 25-cm square piece of 5-mm thick glass floor. To ensure consistent illumination from below, two independent 850-nm near-infrared (NIR) LED strips (SMD5050-300-IR, Huake Light Electronics Co., Ltd., Shenzhen, China) were mounted 10 cm beneath the glass floor. These were carefully positioned to prevent reflections from interfering with the recorded images. The LED strips were powered by a 12-V DC power supply. Video recordings were acquired in darkness using an NIR camera (Basler acA2000-50gmNIR GigE, Basler AG, Ahrensburg, Germany) positioned 30 cm below the glass surface. Pylon viewer software (v4.1.0.3660, Basler) was used to capture videos under standardized camera settings. All recordings were conducted at 25 Hz with an initial resolution of 1000 × 1000 pixels, which was subsequently downsized to 500 × 500 pixels for high throughput analysis. For each injury type, four male and four female C57BL/6J mice were used. To minimize stress and ensure natural behaviour, mice were habituated for 30 minutes in the recording chamber followed by a 30-minute recording session at each time point (Extended Data Fig. 1c,d). Only one mouse was placed in the chamber at a time.

To analyse behaviours from the recordings, we used BAREfoot, a supervised machine learning algorithm that integrates pose estimation with light-based measurements of body-part pressure and distance^18^. For behaviours without pre-existing classifiers, training datasets were generated by manual annotation of videos by experienced scorers, and new classifiers were subsequently trained. The resulting library of trained classifiers enabled automated, frame-by-frame scoring of a behavioural repertoire that included spontaneous rearing, grooming, scratching, biting/licking, guarding, and limb flicks in freely moving mice.

### Home-cage EMG/video recording system

Implantation of wireless telemetry transmitters and recording was performed aseptically in 8-week-old C57BL/6J mice under 2% inhaled isoflurane anaesthesia. Wireless telemetry transmitters (PhysioTel ETA-F10; Data Sciences International, St. Paul, MN) were implanted by making a short skin incision along the dorsal aspect of the left hindlimb. A subcutaneous pocket was created by blunt dissection over the back, cephalad to the hindlimb, to house the transmitter body. For EMG recording, a pair of recording and reference electrode leads were stripped of silicone insulation over a 2–3 mm segment, inserted into the vastus lateralis muscle, and secured to the muscle surface with 5-0 nylon sutures to prevent displacement. The incision was closed with surgical staples and tissue adhesive, and mice were returned to their home cages to recover for ∼2 weeks before undergoing the SNT procedure. For experiments shown in Fig. 1d,f, sham controls included 3 males and 2 females, and the SNT group included 6 males and 4 females. Mice in the sham group were also used to assess weight bearing and gait parameters post-implantation (Extended Data Fig. 3d) using a previously reported behavioural platform ^53^. In a separate cohort, 3 high-pain and 3 low-pain mice at 8 weeks post-SNT were implanted with transmitters to evaluate the impact of implantation on neuroma pain sensitivity (Extended Data Fig. 3e).

Mice were housed in custom-designed home cages equipped with overhead video cameras with night-vision, maintained under a 12-hour light/12-hour dark cycle (lights on at 7 a.m. and off at 7 p.m.) in temperature- and humidity-controlled chambers, with ad libitum access to food and water (Fig. 1a). In a subset of animals, four additional side-view IP cameras (M1145-L Network Camera, Axis) were used. Video/EMG recordings (sampled at 1000 Hz) were acquired in real time using the DSI Ponemah data acquisition platform (Data Sciences International, St. Paul, MN).

To examine the frequency-domain characteristics of EMG signals representing different behaviours, we performed fast Fourier transform (FFT) analysis on raw EMG time series data with Python’s scipy.fft module to determine dominant frequency for each behaviour defined by video observations (Extended Data Fig. 3a). The raw amplitude values of the corresponding time series were first transformed into the frequency domain. The resulting spectrum was computed over the range of 0 to 200 Hz. To facilitate comparison across time series and to highlight relative peak frequencies independent of absolute amplitude differences, the FFT magnitude spectrum for each trace was normalized by dividing all frequency amplitudes by the maximum amplitude within that trace. This rescaled the values to a range between 0 and 1, allowing identification of the dominant frequency peaks while minimizing inter-sample variability in absolute signal strength (Extended Data Fig. 3c, right). Event amplitudes were defined as the absolute peak-to-peak value during each event (Extended Data Fig. 3b). Autocorrelation was computed using the autocorrelation function up to 1000 ms using FFT-based methods. The first positive peak after lag 0 was identified to extract both the dominant periodicity and its autocorrelation strength (Extended Data Fig. 3b).

For scoring limb flicks, raw EMG signals were bandpass filtered at 5–30 Hz. Limb flicks were defined as bouts of repetitive spikes in the filtered EMG trace (Fig. 1a and Supplementary Video 1). Based on features distinguishing limb flicks from other behaviours (Extended Data Fig. 3a,b), we implemented a two-step semi-manual scoring pipeline. First, EMG traces were screened using a dynamic threshold–based automated spike detection algorithm in NeuroScore CNS Data Analysis Software (v3.6; DSI, Data Sciences International, St. Paul, MN) with broad parameters as follows. Threshold ratio: 1.2; Maximum ratio: 3; Minimum value: 5 μV; minimum spike duration: 1 ms; Maximum spike duration: 1000 ms; Minimum inter-spike interval: 10 ms; Maximum inter-spike interval: 1000 ms; Minimum spike train duration: 10 ms; and Minimum number of spikes: 2. This automated step detected >95% of true-positive events as confirmed in pilot observations. Second, automatically detected events were reviewed by an experimenter blinded to the experimental condition (sham vs. SNT and time point post-surgery), who cross-referenced detection timestamps with simultaneous video recordings to exclude false positives. The coefficient of variation was calculated as the standard deviation divided by the mean of flick counts within each binned time window. Ten 24-hour recordings of SNT mice were randomly selected to compute variation across bin sizes ranging from 0.5 to 12 hours. A subset of five mice was recorded for 6 consecutive days to compute CV across extended 12-, 24-, and 48-hour windows (Fig. 1b).

### Transcutaneous neuroma poke test

Mice were shaved around the thigh area before behavioural assay. Acclimation was conducted over two consecutive days prior to the actual test. During acclimation, mice were placed on a laboratory bench while the investigator gently held their tails. A series of pokes (20–50 repetitive trials covering the entire thigh area, ∼15 minutes per mouse) were applied using a 15 g von Frey filament (Stoelting Co, #58011) to familiarize the animals with the testing environment. This acclimation step was crucial, as mice otherwise exhibited escaping behaviour in response to mechanical stimulation, even in the absence of injury. Following acclimation, mice typically remained calm and stationary, indicating readiness for the actual behavioural test (Supplementary Video 2). If mice continued to display escaping or excessive movements, acclimation was extended until they reached a stable state. Additionally, trials during this acclimation process enabled the investigators to identify a ‘target point’—the area on the thigh that elicited the most consistent flicking response. This point often corresponded to the precise location of the neuroma, as later confirmed by *post-hoc* dissection. In mice that did not exhibit a flicking response throughout acclimation, the ‘target point’ was set on the skin area directly above the site of the nerve transection.

For the behavioural test, 10 trials of mechanical stimulation were applied to the identified ‘target point’ using the 15 g von Frey filament, with an inter-stimulus interval of >5 seconds. If a stimulus did not evoke an immediate flicking response, it was maintained for 3 seconds. Trials with no response after this duration were counted as non-responsive (Supplementary Video 2). Since injured mice sporadically exhibited spontaneous flicking behaviours, any response occurring more than 2 seconds after stimulus offset was not counted, and the trial was repeated. Occasionally, mice exhibited other behaviours to pokes (e.g., licking or guarding), which were not quantified as they were rare and also occurred on the contralateral side. Flicking probability to poke was quantified as the proportion of trials that elicited flicking behaviours relative to the total number of trials. For animals tested repeatedly on different days, investigators were blinded to their previously assessed pain phenotype.

### Plantar von Frey assay

Mice were placed individually in transparent plastic chambers on a wire mesh grid and acclimated to the testing environment for 60 minutes on two consecutive days. Calibrated von Frey filaments (0.008–4.0 g) were applied to the medial plantar surface (saphenous nerve territory) of the injured side using the Up-Down method^54^. Testing was performed 1 day before SNT and weekly thereafter up to 8 weeks post-SNT, and 50% withdrawal thresholds were calculated (Fig. 3a).

For plantar von Frey testing in Piezo2 conditional knockout mice generated from *Cdx2^Cre^* (JAX #009350)^55^, *Piezo2^flox^*(JAX #027720) and *Piezo2^flox/Null^* strains^56^, the contralateral central plantar area was targeted using the same Up-Down method (Extended Data Fig. 5).

### Drug treatments and chemogenetic silencing experiments

For the drug-related tests, spontaneous behavioural assessments were performed in the black box recording system (Extended Data Fig. 1b) for 30 minutes rather than in the home-cage monitoring setup, as the effective window of drug action is on the scale of hours. To mitigate intra-animal variability and ensure robust statistical power, a larger sample size was included. Gabapentin (Medisca, #60142-96-3) was administered intraperitoneally at 30 mg/kg. Spontaneous behaviours were recorded in the black box for 30 minutes before injection (pre) and again for 30–60 minutes post-injection (Extended Data Fig. 2a). Compound 21 (Hello Bio, #HB4888) was administered intraperitoneally at 3 mg/kg to mice carrying either the hM4Di allele (B6.129-*Gt(ROSA)26Sor^tm1(CAG-CHRM4,-mCitrine)Ute^*/J; JAX #026219) alone, *Sstr2^T2a-CreER^* alone, or hM4Di under the control of *Sstr2^T2a-CreER^*. Transcutaneous neuroma poke tests were conducted 40–50 minutes post-injection (Fig. 4f). 50 µl 1% lidocaine hydrochloride monohydrate (Sigma-Aldrich, #L5647) was administered via deep intramuscular–perineural injection around the lateral upper thigh where the neuroma resides. Spontaneous behaviours were recorded for 30 minutes beginning 15 minutes after injection, immediately followed by transcutaneous neuroma poke tests conducted 45–55 minutes post-injection (Fig. 1h). For calcium imaging experiments, 10 µL of 1% lidocaine hydrochloride monohydrate was applied either topically to surgically exposed neuromas (Fig. 2h and Fig. 3g) or via intraplantar injection (Fig. 3h), and von Frey assessments were performed 5 minutes after administration. Capsaicin (Sigma-Aldrich, #M2028), zymosan (Sigma-Aldrich, #Z4250), and CFA (Sigma-Aldrich, #F5881) were administered as described in the legend to Extended Data Fig. 2b. Spontaneous limb flicks and paw biting/licking behaviours were manually counted by investigators blinded to the treatments.

### *In vivo* imaging of DRG activity

DRG imaging was performed as described previously^38^. Mice carrying both *Vglut2-ires-Cre* (B6J.129S6(FVB)-*Slc17a6^tm2(cre)Lowl^*/MwarJ; JAX #28863) and *Ai95D* (B6J.Cg-*Gt(ROSA)26Sor^tm95.1(CAG-GCaMP6f)Hze^*/MwarJ; JAX #028865) alleles were used for imaging at 8–12 weeks post-SNT. Anaesthesia was induced with 2.5% isoflurane and maintained at 2% via a nose cone. Mice were placed on a heated pad to maintain the body temperature at 37 °C, and ophthalmic ointment was applied to the eyes throughout the surgery and imaging. The back was shaved, and a rectangular incision was made between the lumbar enlargement and pelvic bone with the skin folded away. A dorsal laminectomy was performed in four steps. First, a 1.5-cm midline incision caudal to the lumbar enlargement area was made to create space for clamping. Second, a 3–4 mm incision was made approximately 1 cm rostral to the pelvic bone to expose the spine where the L4 DRG is situated. Third, muscles and connective tissues were removed with scissors and forceps. Fourth, the L4 transverse process was excised using rongeurs, and small vertebral fragments were carefully trimmed with fine forceps to fully expose the DRG. Bleeding was controlled using absorbent paper points (Electron Microscopy Sciences, #71011-01). To minimize motion artifacts from respiration, a custom spinal clamp was used to secure the spine over the vertebrae 5 mm rostral to the L4 DRG. Illumination was provided with a collimated 470 nm light-emitting diode through a 10x, 0.3 numerical aperture air objective. The clamp was adjusted to ensure the DRG surface was as level as possible, visualizing the maximum surface area in the focal plane, typically covering ∼80% of the entire DRG. A single focal plane was selected to maximize the number of clearly resolvable neurons across the DRG exhibiting robust calcium responses to indentation of the medial hindpaw (saphenous nerve territory). A 30-minute imaging session without external stimulation was conducted to assess spontaneous activity. The neuroma was then surgically exposed following a procedure similar to sciatic nerve exposure during SNT injury. Mechanical stimuli of varying forces were applied to the neuroma using von Frey filaments. Each force was delivered in five trials, with an interstimulus interval of >5 seconds. After imaging the ipsilateral L4 DRG, the same procedure was repeated for the contralateral L4 DRG, except that mechanical stimuli were applied to the intact sciatic nerve at a location corresponding to the neuroma site on the ipsilateral side. The imaging order (i.e., ipsilateral or contralateral first) was randomized to minimize potential variability due to anaesthesia duration. Images were acquired at 10 Hz with a 99 ms exposure time during neuroma stimulation. The stimulation was also recorded simultaneously at 25–30 Hz using a digital camera, with the video output time-stamped to synchronize with GCaMP imaging data. For imaging spontaneous activity, exposure time was reduced to 35 ms to minimize photobleaching during long-term recordings. Isoflurane concentration was reduced to 1% throughout the imaging process.

Imaging data were analysed using the open-source CaImAn package^57^. Briefly, data were first corrected for motion using the NoRMCorre algorithm. Regions of interest (ROIs) were identified using a constrained non-negative matrix factorization (CNMF) framework, which jointly estimates spatial footprints and temporal activity traces. Extracted fluorescence traces were detrended to correct for bleaching and converted to ΔF/F on a per-cell basis, with baseline (F) defined as the average of the lowest 10% of fluorescence values. Automatically extracted ROIs were reviewed by an investigator to correct false positives and false negatives. The corresponding calcium traces were then evaluated against quality criteria, including spatial correlation, temporal signal-to-noise ratio (ΔF/F exceeding 3× the standard deviation of baseline fluctuation for ≥0.5 s), appropriate rise and decay kinetics, and consistent morphology. Only ROIs meeting these criteria were retained for analysis.

To classify spontaneously active cells, neuroma poke–responsive cells, and dual-active cells based on calcium activity, we used an unsupervised clustering approach with Gaussian Mixture Models (GMMs) implemented in Python’s scikit-learn package. The model estimated the probability of each cell belonging to one of three classes (Fig. 2g) or two classes (Fig. 3d) according to its peak ΔF/F across the corresponding imaging sessions, with final cluster assignment determined by maximum posterior probability (decision boundary).

### Miniaturized LED cuff and epineural optogenetic activation

We adapted a previously developed wireless LED array^32^ into a wired device to enable behavioural screening across multiple Cre lines (Fig. 4b and Extended Data Fig. 7). The fabrication process for the LED cuff began with the spin-coating of a 3 μm-thick polyimide (PI) layer (HD Microsystems GmbH, catalog no. PI2611) onto a silicon (Si) wafer coated with an aluminum (Al) layer. The underlying Al acted as a sacrificial layer for subsequent device release. The PI was then baked at 300 °C for 2 hours to complete the imidization process. A 130 nm-thick silicon carbide (SiC) layer was then deposited via plasma-enhanced chemical vapor deposition (PECVD) using an Oxford PlasmaLab System 100 under the following conditions: 133 sccm Ar, 60 sccm CH₄, and 375 sccm of 2% SiH₄ in Ar; RF power of 25 W; substrate temperature of 300 °C; and deposition time of 10 minutes. Next, a metal stack composed of Titanium/Platinum/Gold/Platinum/Titanium (Ti/Pt/Au/Pt/Ti, thicknesses: 10/10/200/10/10 nm) was sputtered using an AC450 system (Alliance-Concept) and patterned by lift-off in an ultrasonic acetone bath (80 kHz, 100% power, 10 minutes). A second 130 nm SiC layer was deposited, followed by spin-coating and baking of an additional PI layer using the same parameters as before. A Titanium/Silicon Oxides (Ti/SiO₂) bilayer (10/25 nm) was subsequently sputtered onto the top PI surface. The final device patterning was carried out using an ICP-RIE etch system (Corial 210IL), defining the shape of the cuff and opening windows for LED integration and soldering. Following the microfabrication steps, solder paste bumps (∼50 μm diameter, Chipquik, catalog no. SMDLTLFP10T5) were manually deposited onto the exposed contact pads. Micro-LEDs (Cree Inc., catalog no. DA2432) were placed using a pick-and-place system (JFP Microtechnic) and bonded by heating the wafer on a hotplate at 165 °C, ensuring both electrical and mechanical connections. To improve hermetic sealing, a 15 wt% solution of poly(isobutylene) (PIB, Oppanol, BASF) in cyclohexane (Sigma-Aldrich) was applied to the die surface via pneumatic dispensing. Finally, a 60 μm-thick encapsulating layer of polydimethylsiloxane (PDMS) was printed, baked at 70 °C for 12 hours, and patterned by blade cutting. DC voltage was measured using a source meter (Keithley 2400) while sweeping the input current from 0.5 to 17 mA. Simultaneously, the total optical power output at 470 nm was measured using a calibrated photodiode (S170C, Thorlabs) connected to a power meter console (PM100D, Thorlabs). During measurements, the distance between the photodiode and the micro-LEDs was maintained at approximately 1 mm.

The neuroma or intact sciatic nerve was exposed as described in the sciatic nerve injury procedure. The LED cuff was positioned beneath the neuroma or intact sciatic nerve at a perpendicular angle, and its distal end was folded 180° to wrap around the target site. This configuration placed one set of LEDs on the front and the other on the back of the neuroma or nerve (Fig. 4b, right). To secure the device, the two proximal anchoring points were sutured to each other and to the adjacent proximal muscles, while the distal anchoring points were sutured to the upper thigh muscle for additional stabilization (Fig. 4b, right). 10-0 sutures (Ethicon, #2794G) were used for these anchoring sites. The wires with a female nano-miniature circular connector (Omnetics, #A79103-001) were routed subcutaneously to the backside, where a small skin pocket was created to externalize the connector. The thigh incision was closed using 6-0 sutures, while the skin pocket was sealed with tissue adhesive (3M Vetbond, #1469). After each experiment, the placement of the LEDs was confirmed. If displacement occurred (e.g., slippage from the neuroma), the animal was excluded from analysis.

The following transgenic lines were used to target sensory neuron subtypes: *Sstr2^T2a-CreER^;Calca-FlpE;R26^LSL-FSF-ReaChR::mCitrine^*for the Neuroma C-Nociceptor (SSTR2) subtype, *Mrgprd^CreER^;R26^LSL-ReaChR::mCitrine^*for the C-HTMR/HEAT (MRGPRD) subtype, *Th^T2a-CreER^;Avil^FlpO^;R26^LSL-FSF-ReaChR::mCitrine^*for the C-LTMR subtype, *Smr2^T2a-Cre^;Calca-FlpE;R26^LSL-FSF-ReaChR::mCitrine^* for the Aδ-HTMR/Heat (SMR2) subtype, *TrkB^CreER^;Avil^FlpO^; R26^LSL-FSF-ReaChR::mCitrine^* for the Aδ-LTMR subtype, *Mrgpra3^Cre^;R26^LSL-ReaChR::mCitrine^* for the C-HTMR/Heat (MRGPRA3) subtype, *Mrgprb4^Cre^;R26^LSL-ReaChR::mCitrine^*for the C-HTMR/Heat (MRGPRB4) subtype, *Cysltr2^Cre^;R26^LSL-ReaChR::mCitrine^* for the C-HTMR/Heat (CYSLTR2/SST) subtype, *Trpm8^T2a-FlpO^; R26^FSF-ReaChR::mCitrine^* for the C-Cold (TRPM8) subtype. All the mouse lines used to generate these genetic crosses have been previously described: *Sstr2^T2a-CreER^* (JAX #039563)^28^, *Mrgprd^CreER^* (JAX #031286)^58^, *Th^T2a-CreER^* (JAX #025614)^59^, *Smr2^T2a-Cre^* (JAX #039562)^28^, *TrkB^CreER^* (JAX #027214)^60^, *Mrgpra3^Cre61^*, *Mrgprb4^Cre^*^62^, *Cysltr2^Cre^* (JAX #039985)^28^, *Trpm8^T2a-FlpO^* (JAX #039564^28^, *Calca-FlpE^63^*, *Avil^FlpO63^*, *R26^LSL-FSF-ReaChR::mCitrine^* (JAX #024846), *R26^LSL-ReaChR::mCitrine^* (JAX #026294), *R26^FSF-ReaChR::mCitrine^* (derived from JAX #024846 by germline excision of LSL cassette). All transgenic mouse lines were maintained on a mixed background and included males and females. CreER lines were induced by a single intraperitoneal injection of tamoxifen dissolved in sunflower seed oil, prepared as previously described^28^. For *Sstr2^T2a-CreER^; Calca-FlpE; R26^LSL-FSF-ReaChR::mCitrine^*, we administered 3 mg of tamoxifen at 3–4 weeks; *Mrgprd_CreER_;R26_LSL-ReaChR::mCitrine_*, 2 mg at 3-4 weeks; *Th_T2a-CreER;_Avil_FlpO_;R26_LSL-FSF-ReaChR::mCitrine_*, 2 mg at 3-6 weeks; *TrkB^CreER^; Avil^FlpO^; R26^LSL-FSF-ReaChR::mCitrine^*, 0.5 mg at postnatal day 3 to day 5. For optogenetic activation, the external connector was coupled to an adapter linked to a stimulator (AM-Systems, Grass Instruments S88). When mice were in a calm, awake state, a single 10 ms pulse was delivered at various voltages corresponding to different optical power levels with behaviours recorded by a high-speed camera at 120 Hz. Each power level was tested in 10 trials with an inter-stimulus interval of >1 minute. Behaviours occurring during stimulation or within 2 s after stimulus offset were counted as positive responses. Response probability was calculated as the proportion of trials with a behavioural response out of 10 trials for each animal (Fig. 4c). The occurrence rate of each behaviour type was calculated as the number of trials in which that behaviour was induced from the ipsilateral neuroma site at 60 mW across all animals, divided by the total number of trials (i.e., 10 × number of animals) (Fig. 4d). Response latency (Fig. 4e, left) was defined as the time between opto-stimulation onset (single 10 ms pulse at 60 mW) and the initiation of the behaviour. Single 10 ms pulses were used for all optogenetic experiments except in Extended Data Fig. 9a, where repetitive 10 ms pulses at 10 Hz or continuous light (60 mW) were delivered for 5 s. Cre expression in all animals was verified *post hoc* by histological analysis following completion of optogenetic experiments (Extended Data Fig. 8).

Survival analysis using a Cox proportional hazards regression model was used to assess whether flick probability to transcutaneous neuroma pokes predicted the optical threshold for evoked limb flicks. During optogenetic stimulation, optical power (mW) was increased monotonically and treated as the ordered analysis variable, with the first limb flick defined as the event. Animals that did not flick at the maximum tested power (60 mW) were right-censored. Von Frey flick probability (0–1) was included as a continuous covariate. Hazard ratios with 95% confidence intervals were reported, and significance was assessed using the likelihood-ratio test (Fig. 4e, right).

### Histology

Neuromas, contralateral intact sciatic nerves, or DRG were fixed in 4% paraformaldehyde (PFA) in PBS at 4 °C overnight. Nerves were cryoprotected in 30% sucrose in PBS at 4 °C for one to two days, embedded in OCT compound (Sakura Finetek) on dry ice, and cryosectioned longitudinally (parallel to the axon trajectory) at 50 µm. DRG were fixed and processed for whole-mount staining. Sections or whole DRG were permeabilized in 1% Triton X-100 in PBS for 20 minutes and blocked in 10% normal donkey with 0.3% Triton X-100 in PBS overnight at 4 °C. Primary antibody incubation was performed overnight at 4 °C in blocking solution (1:500) using anti-GFP (Thermo Fisher, #A10262) and anti-PGP9.5 (Abcam, #ab108986). Secondary antibodies were incubated for 2 hours at room temperature or overnight at 4 °C in blocking solution (1:500): Cy5-conjugated donkey anti-rabbit (Jackson ImmunoResearch, #711-175-152) and Alexa Fluor 488–conjugated donkey anti-chicken (Jackson ImmunoResearch, #703-545-155). Images were acquired on a Leica SP8 confocal microscope at 1024 × 1024 resolution. Whole DRG were imaged in 60 mm glass-bottom dishes using 10× or 20× air objectives with 2–5 µm z-steps. Neuromas and contralateral nerves were captured by tile scan using a 10× air objective with 2 µm z-steps. Post-processing was performed with LAS X Life Science software (Leica).

fibre density (Extended Data Fig. 8) was quantified from mCitrine reporter signal in neuromas and contralateral sciatic nerves using ImageJ. Each neuroma and contralateral nerve typically yielded ∼20 and ∼12 longitudinal cryosections, respectively; the middle three consecutive slices were selected to minimize epineural tissue interference. Images were converted to 8-bit grayscale, background-subtracted, and thresholded to generate binary masks of axonal fibres. A 50 × 50 µm region of interest (ROI) was placed at the neuroma tip or corresponding contralateral nerve center, and the axon-occupied area within this ROI was measured. fibre density was expressed as the fraction of axon-positive area relative to total ROI area. Values from three slices were averaged to obtain one measurement per sample.

### *In situ* hybridization combined with immunohistochemistry

L4 DRG were dissected under 2% isoflurane anaesthesia and embedded in OCT on dry ice. The tissues were then cryosectioned at a thickness of 20 µm and stored at -80°C until processing. To assess the co-expression of *Sstr2* mRNA and mCitrine protein (Extended Data Fig. 9b), *in situ* hybridization (ISH) was performed using the RNAscope (Advanced Cell Diagnostics) combined with immunohistochemistry (IHC). Sections were processed according to the manufacturer’s ISH instructions using the RNAscope H2O2 and Protease Reagents, the RNAscope Multiplex Fluorescent Detection Kit v2, with *Sstr2* (Cat. # 437681-C2) detected via Akoya Opal Dye 570.

Immediately following the RNAscope protocol, IHC was performed to detect mCitrine expression. Briefly, slides were blocked with 10% Normal Donkey Serum with PBST (1X PBS with 0.1% Tween20) for 15 minutes at room temperature. Primary antibody incubation was performed for 1 hr at room temperature in blocking solution using goat anti-GFP antibody (1:500; US Biological Life Sciences, G8965-01E). Following two 5-minute rinses in PBST, secondary antibodies incubation was performed for 40 min at room temperature in blocking solution using Donkey anti-Goat Alexa Fluor 488 (ThermoFisher Scientific, A-11055). After the final washes, sections were counterstained with DAPI and coverslipped. Confocal images were acquired using a Zeiss LSM 900 microscope equipped with 10X and 20X air objectives.

### snRNA-seq of DRG across pain conditions

From a parent cohort of 60 mice, L4 and L5 DRG were collected from 18 mice (9 males and 9 females) that consistently exhibited a high-pain phenotype across 8–10 weeks post-SNT, with phenotyping performed weekly and tissue collected at 10 weeks, and from 18 mice (9 males and 9 females) that consistently exhibited a low-pain phenotype over the same period. 36 contralateral L4 and L5 DRG from both high- and low-pain mice were combined as the control group.

All nuclear extraction steps were performed on ice, or at 4 °C. Frozen L4 and L5 DRG, collected on dry ice in DNA/RNA LoBind tubes (Qiagen) were homogenized in NST buffer: 50 ml NST buffer contained 25 ml of 2x ST buffer, 1ml of 10% IGPAL, 1.4 ml of 35% BSA, and 22.6 ml nuclease-free water, and was prepared from ST buffer, containing 146 mM NaCl, 10 mM Tris-HCl, pH 7.5, 1 mM CaCl_2_, and 21 mM MgCl_2_^64^. For each sample, 7.5 ml of ST and 7.5 ml of NST buffer were prepared and 10 µl of Rnasin was added to each. For each sample, 1 ml cold NST buffer (on ice) was added directly to the tube containing frozen DRG, allowed to thaw for 15–30 seconds, and DRG were transferred to a 7 ml Dounce Wheaton homogenizer containing 4 ml of NST buffer, and homogenized with 5 strokes with the loose pestle and 15 strokes with the tight pestle. The homogenate was then filtered through a 40 µm cell filter into a 50 ml Falcon tube on ice. 5 ml of ST buffer was added to the homogenizer, and any remaining homogenate was collected and also pipetted through the 40 µm cell filter. 10 ml of filtered homogenate then filtered through a 20 µm cell filter into a 15 ml Falcon tube on ice. The homogenizer was thoroughly rinsed between samples. Samples were centrifuged at 500 × g for 10 minutes at 4 °C. The supernatant was removed and the pellet was resuspended in 1 ml of ST buffer. 10 µl of Hoechst was added for a final concentration of 0.5 µg/ml, and the samples were FACS sorted to isolate nuclei.

Nuclei suspensions were sequenced using 10x Genomics assays, resuspended, and loaded into the 10x Chromium device for snRNA-seq (10x Genomics 3’ v3). Libraries were prepared for snRNA-seq according to the manufacturer’s protocol. Libraries were sequenced on an Illumina NovaSeqX (Azenta). Sequencing data were processed and mapped to the mouse (mm10) genome using 10x Genomics cellranger v9 using default parameters. We ran the remove-background function of Cellbender (v0.3.0) to remove ambient RNA and empty droplets from raw expression matrices using default parameters.

We followed a similar cell type annotation and clustering approach as previously described^30,47^, and adopted the established physiological nomenclature for each neuronal subtype as shown in Table 1 in a reference study^28^. Seurat v4 was used to perform quality control, data normalization, cell type annotation and comparison between neuroma conditions^65^. Nuclei with >1000 genes and <10% of mitochondrial counts were included for analysis. We found 2000 variable genes for each library after log-normalization, and then integration anchors were selected across all libraries. The data were then scaled, and we selected PCs on the basis of the point where the PC only contributes to 5% of SD of highly variable gene expression and all PCs cumulatively contributed to 90% of the SD of highly variable gene expression. The libraries were then integrated using Seurat’s CCA algorithm, and nuclei were clustered and plotted in UMAP space at a resolution of 0.5. A Wilcoxon rank sum test was performed between the clusters to identify marker genes. We annotated clusters that were *Snap25^+^* (log_2_FC > 0.5 and adjusted *P* value < 0.05 relative to all other clusters) as “neurons” and clusters that were *Sparc*^+^ (log_2_FC > 0.5 and adjusted *P* value < 0.05 relative to all other clusters) as “non-neurons.” Neurons were then separated and reintegrated using a resolution of 1. We annotated neurons using marker gene expression (Extended Data Fig. 6a; log_2_FC > 0.5; adjusted p-value < 0.05 relative to all other clusters). Clusters were labelled as doublets if they contained both expression of *Snap25*, *Mpz*, and *Sparc* (log_2_FC > 0.5 and adjusted *P* value < 0.05 relative to all other clusters). Doublets were then removed, and the nuclei were reintegrated using the same parameters as described and then reannotated for a final time. To identify genes differentially expressed between conditions within each subtype, we used the Wilcoxon rank-sum test.

Directional concordance scores for each gene within each cell type were computed from its log₂ fold-change (log₂FC) values between the high-pain vs. low-pain and high-pain vs. control comparisons (Extended Data Fig. 6c), defined as:

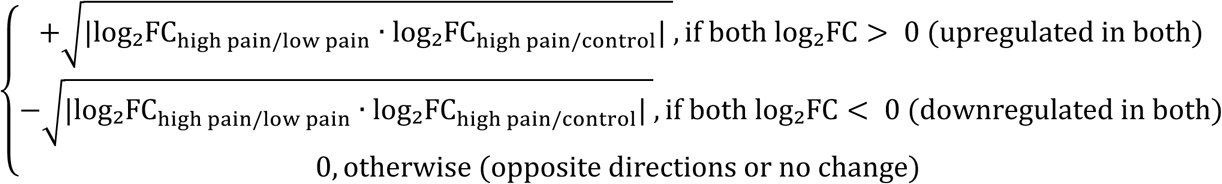

For gene ontology (GO) analysis, we used EnrichR ^66^ to assess functional enrichment. We selected genes significantly enriched in the Neuroma C-Nociceptor (SSTR2) population under the high pain condition (log_2_FC > 0.5; adjusted *p* < 0.05), compared to either the Neuroma C-Nociceptor (SSTR2) cells in the low-pain condition or the contralateral control condition. As the background gene list, we used all genes expressed in the Neuroma C-Nociceptor (SSTR2) population.

### Statistical analysis

All statistical analyses, except for snRNA-seq data, were performed using GraphPad Prism, with the exact tests and *P* values reported in the figures, figure legends, or source data.

## Data availability

All data supporting the findings of this study are available within the manuscript. Reagents are available from the corresponding authors upon reasonable request. snRNA-seq will be deposited into GEO upon manuscript acceptance.

## Acknowledgements

We thank Dr. Bruce P. Bean and members of the Ginty and Woolf laboratories for their constructive feedback. We thank Kitwa Ng for helpful comments on the manuscript. We also thank the IDDRC Animal Behavior and Physiology Core, funded by NIH/NICHD P50HD105351 and IDDRC Cellular Imaging Core, funded by NIH P50 HD105351 at Boston Children’s Hospital. Schematics and illustrations were created with BioRender.com. This work was supported by the International Human Frontier Science Program Organization LT0026/2023-L (X.Z.), NIH grants R35NS105076 (C.J.W.), R35NS097344 (D.D.G.), R35NS132196 (D.D.G.), and R01AT011447 (D.D.G. and C.J.W.), the Edward R. and Anne G. Lefler Center for Neurodegenerative Disorders (D.D.G.), and the K. Lisa Yang Brain Body Center at Harvard Medical School (D.D.G.). D.D.G. is an investigator of the Howard Hughes Medical Institute. This article is subject to HHMI’s Open Access to Publications policy. HHMI lab heads have previously granted a nonexclusive CC BY 4.0 license to the public and a sublicensable license to HHMI in their research articles. Pursuant to those licenses, the author-accepted manuscript of this article can be made freely available under a CC BY 4.0 license immediately upon publication.

## Author contributions

X.Z., D.D.G., and C.J.W. conceived the study. X.Z. performed all experiments and analysed the data, with assistance from R.M.G. (generation of transgenic mouse lines and *in situ* hybridization), S.A.B. (snRNA-seq), K.H. (histology), N.M.B., E.B., E.S. (behavioural studies and histology), K.W. (LED cuff design), O.B. (automated behavioural screen), B.L.T. (acute pain experiments), S.S.J., B.G.E. (tissue collection for snRNA-seq), M.Q.H. (home-cage EMG setup), C.S. (Piezo2-knockout experiments), K.L. (generation of transgenic mouse lines and *in situ* hybridization), E.O.A. (generation of transgenic mouse lines), and I.F. (LED cuff calibration). A.R. provided resource for the home-cage EMG experiments. B.R.J. provided expertise on and assisted with the transcutaneous neuroma poke tests. B.J.W. provided expertise on the behavioural experiments. W.R. provided resource and oversight of the snRNA-seq experiments. S.P.L. provided resource and oversight of LED cuff design. D.D.G. and C.J.W. supervised the project. X.Z., D.D.G., and C.J.W. wrote the manuscript with input from all authors.

---

**Extended Data Fig. 1.**
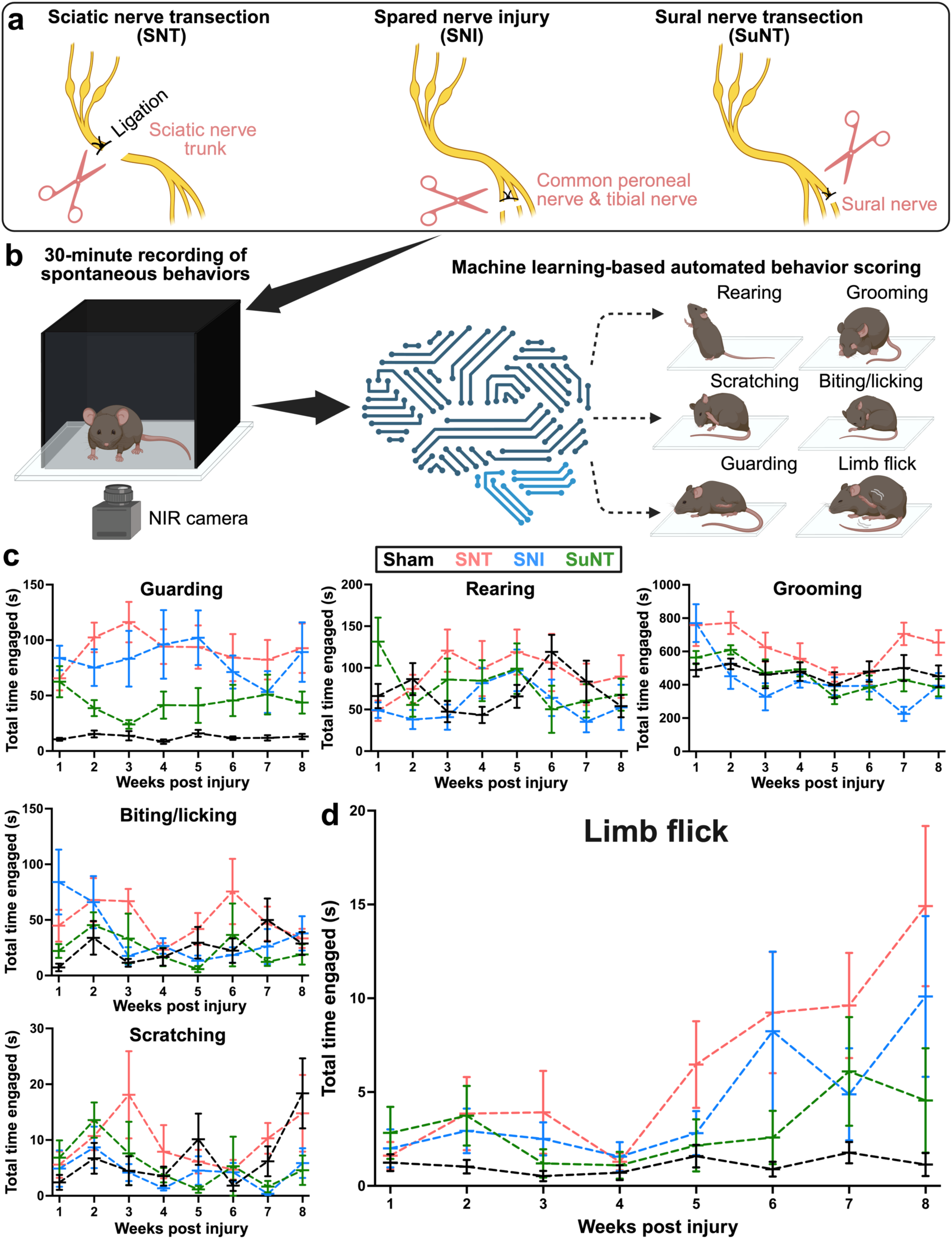
| Automated behavioural screening reveals limb flick as a prominent behaviour in the chronic phase after peripheral nerve injury. **a**, Schematics of three sciatic nerve injury models: sciatic nerve transection (SNT), spared nerve injury (SNI), and sural nerve transection (SuNT). **b**, Experimental setup for 30-minute recordings of spontaneous behaviours using a bottom-up near-infrared (NIR) camera and machine-learning–based automated scoring^18^. **c**,**d**, Total time spent on each indicated behaviour on the injured side, measured weekly for up to 8 weeks post-injury. Lines represent group means, error bars indicate ± SEM. For each injury type, n = 8 mice (4 males, 4 females).

**Extended Data Fig. 2.**
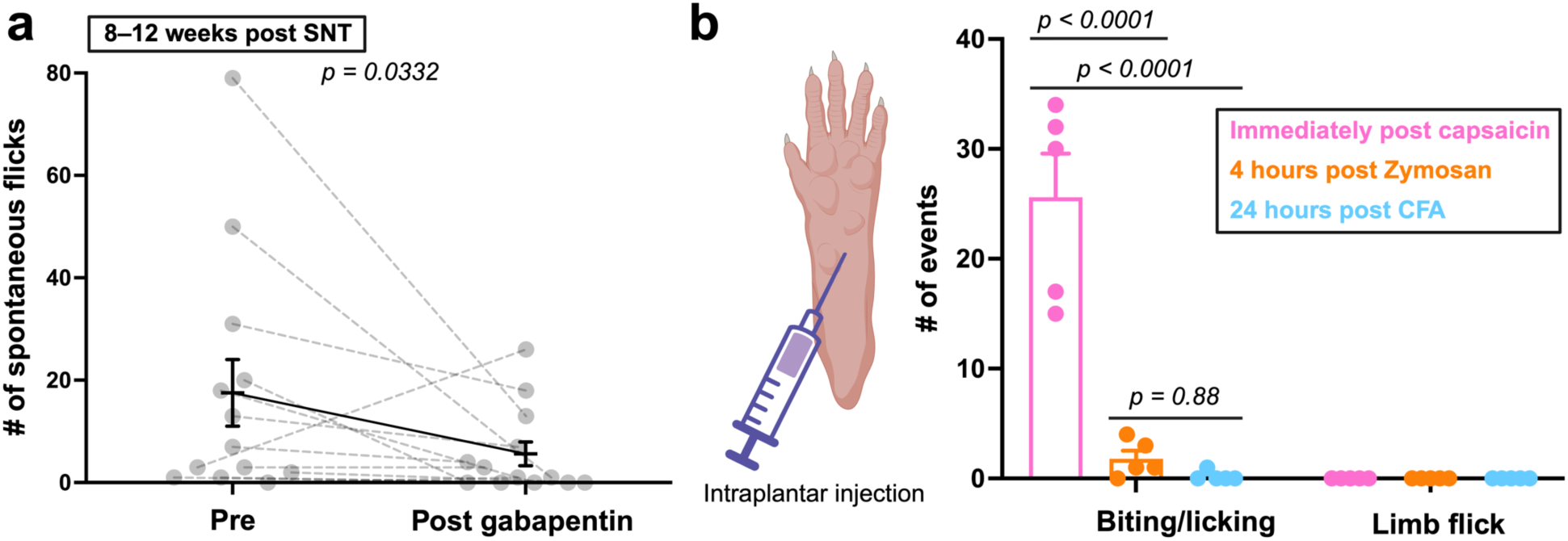
| Limb flicks are suppressed by gabapentin and are absent in acute pain conditions. **a**, Spontaneous limb flicks quantified over 30 minutes before and after intraperitoneal gabapentin (30 mg/kg) at 8–12 weeks post-SNT (n = 13 mice). Data are mean ± SEM; dashed lines connect individual mice. *P* value from two-sided Wilcoxon matched-pairs signed-rank test. **b**, Schematic of intraplantar injection (left) and quantification of biting/licking events and spontaneous limb flicks (right) following capsaicin (0.01%, 10 µL), Zymosan (100 µg in 10 µL), or complete Freund’s adjuvant (CFA, 100%, 10 µL) injection. Behaviours were recorded for 30 minutes at peak-response windows defined from literature: immediately after capsaicin^18^, 4 hours after Zymosan^53^, and 24 hours after CFA^67^. Limb flicks are absent across all acute pain models, while biting and licking of the affected paw is the predominant behaviour in the capsaicin model. Each dot represents one mouse (n = 5 per model). Data are mean ± SEM; *P* values from one-way ANOVA with Tukey’s multiple comparisons.

**Extended Data Fig. 3.**
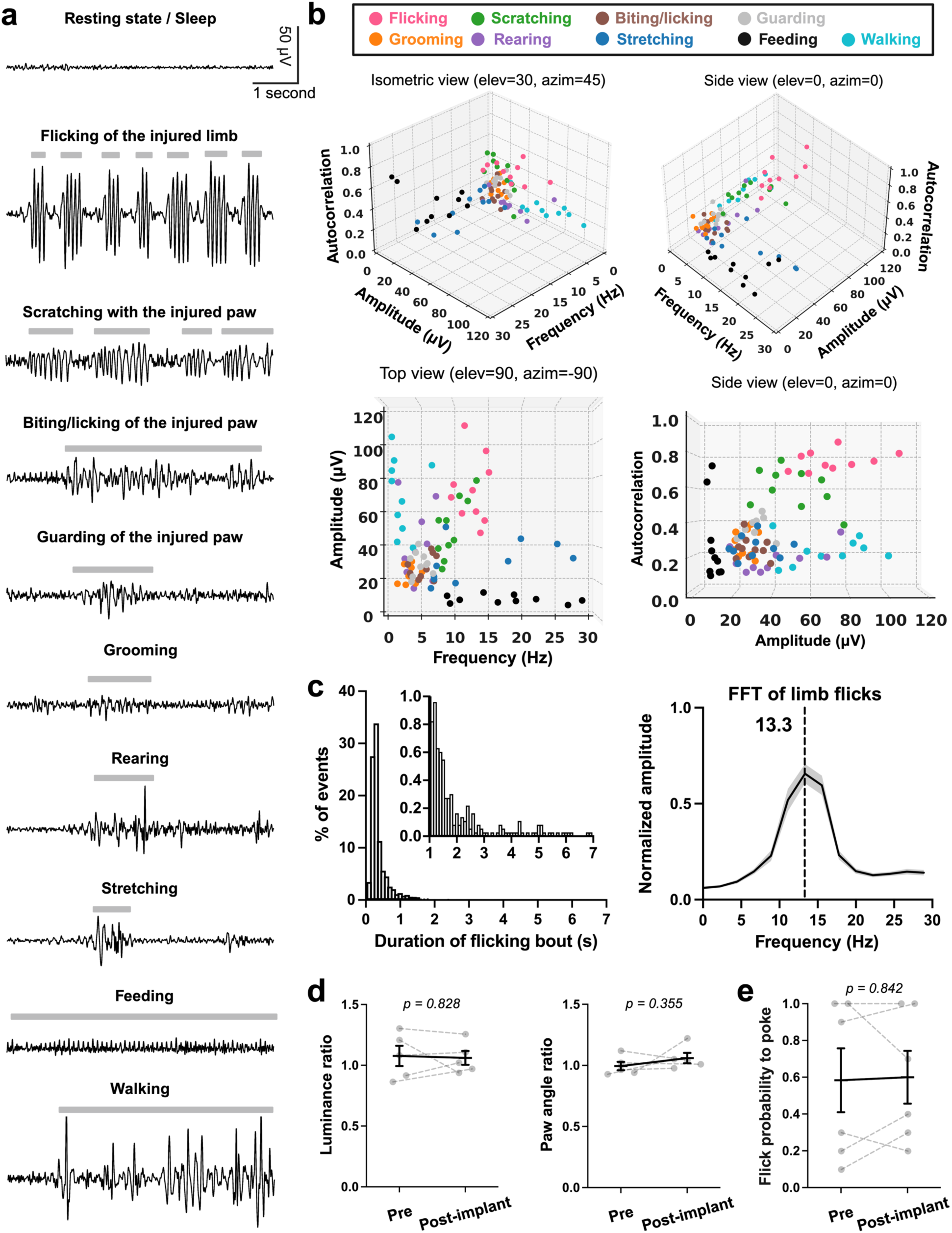
| 24-hour EMG-based home-cage monitoring captures spontaneous limb flicks. **a**, Representative EMG traces during various home-cage behaviours. Gray bars indicate periods when the behaviours occurred as confirmed by corresponding videos. **b**, Feature plots of peak amplitude, frequency, and autocorrelation coefficient (first non-zero positive peak) of EMG trace for each behaviour. Periodic movements such as limb flicks showed higher autocorrelation values. Each dot represents the average of 20 events from one mouse (n = 10 mice, 12 weeks post-SNT). Elev, elevation; azim, azimuth. **c**, Left: histogram showing the duration distribution of individual limb flick bouts, with an inset expanding the 0–1% vs 1–7 seconds range. Most events occurred within 1 s. Right: Fast Fourier Transform (FFT) of limb flicks showing a dominant frequency at 13.3 Hz. **d**, Weight bearing and gait parameters measured by luminance ratio (left) and paw angle ratio (right)^53^ before and two weeks after implantation of the wireless telemetry device without nerve injury (n = 5 mice). **e**, Flick probability to poke before and two weeks after implantation of the wireless EMG device (n = 6 mice at 8–10 weeks post-SNT). Data are mean ± SEM; dashed lines connect individual mice. *P* values from two-sided Wilcoxon matched-pairs signed-rank tests in (**d**,**e**).

**Extended Data Fig. 4.**
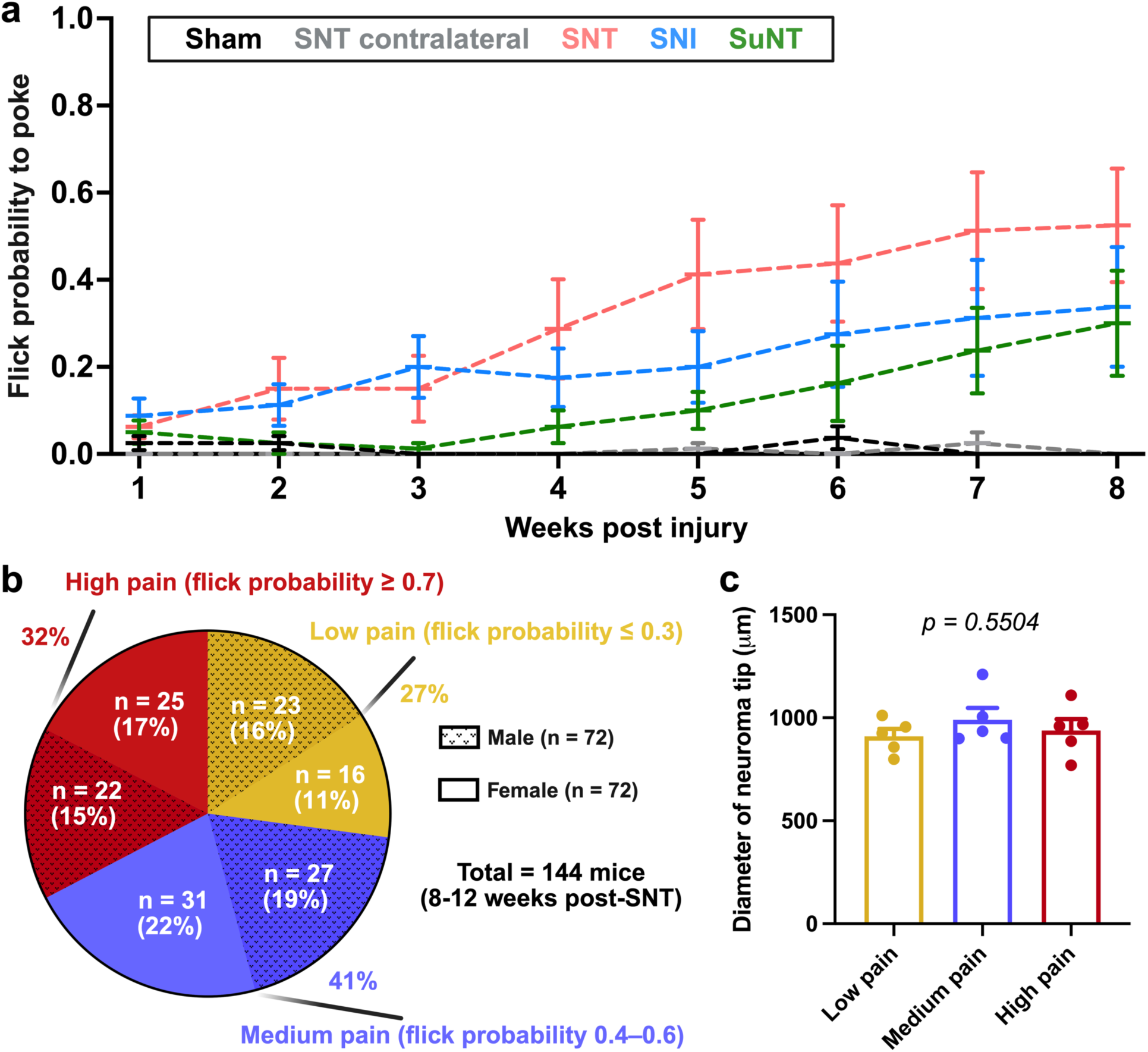
| Neuroma pain emerges across peripheral nerve injury models and shows differential pain severity. **a**, Flick probability in response to transcutaneous neuroma poke measured weekly for up to 8 weeks after injury. Lines represent group means; error bars indicate ± SEM. For each injury type, n = 8 mice (4 males and 4 females, as in Extended Data Fig. 1). **b**, Pie chart showing the distribution of high-pain, low-pain, and medium-pain phenotypes. **c**, Histogram showing the size of neuromas in each pain condition. Each dot represents a neuroma (n = 5 mice for each condition). *P* value from one-way ANOVA.

**Extended Data Fig. 5.**
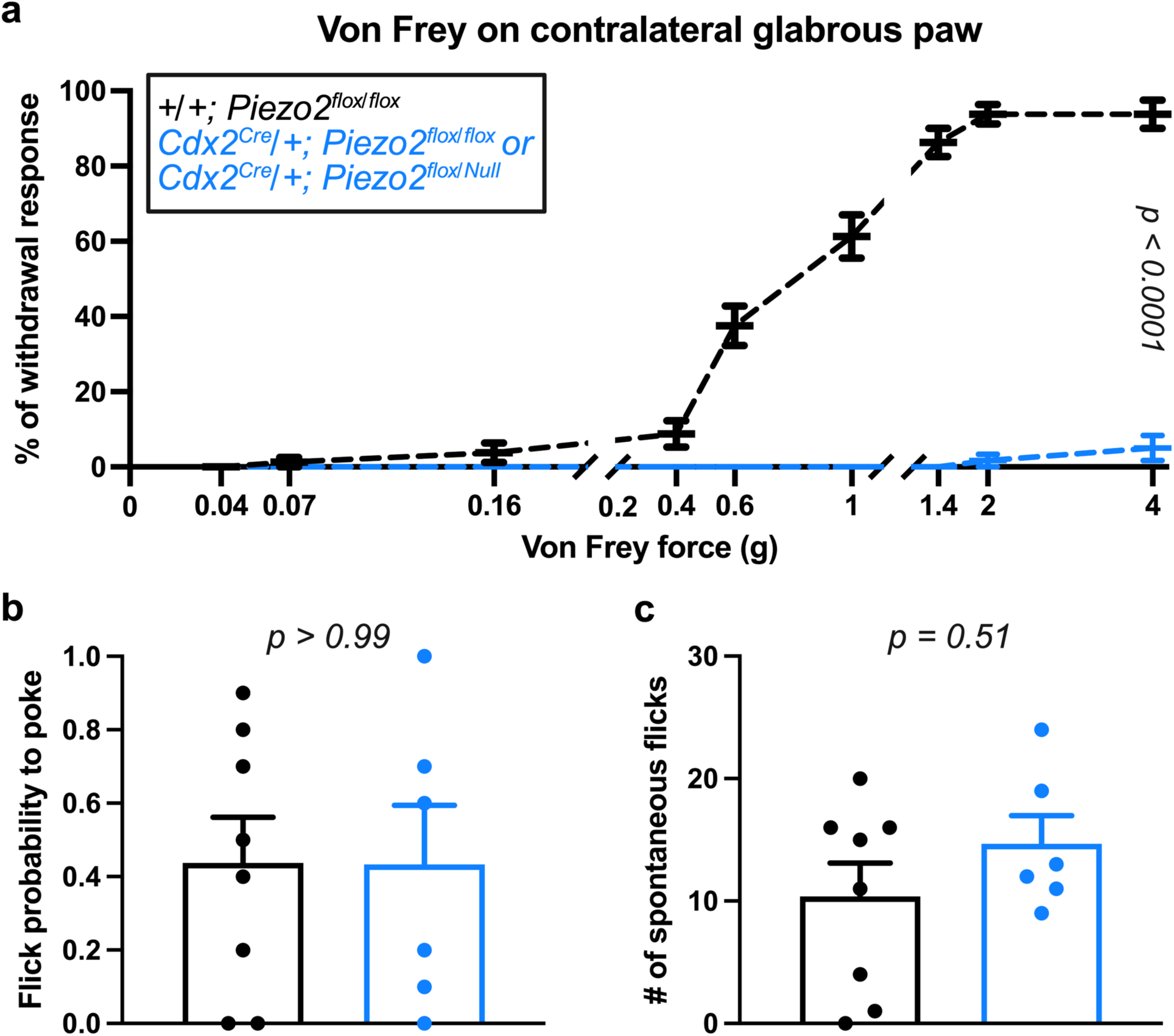
| Spontaneous limb flicks and flicks evoked by transcutaneous poke are independent of Piezo2. **a**-**c**, Von Frey filament evoked responses from contralateral glabrous paw (**a**), flick probability to transcutaneous neuroma poke (**b**), and spontaneous flick count over 30 minutes (**c**) at 8 weeks post-SNT in control and Piezo2 conditional knockout mediated by *Cdx2^Cre^*^55^. Data are mean ± SEM; each dot represents one mouse (n = 8 controls, 6 knockouts). *P* values in (**a**) from two-way ANOVA with Tukey’s multiple comparisons (figure shows the interaction effect of control vs knockout across von Frey forces), and from two-sided Mann–Whitney tests in (**b**,**c**).

**Extended Data Fig. 6.**
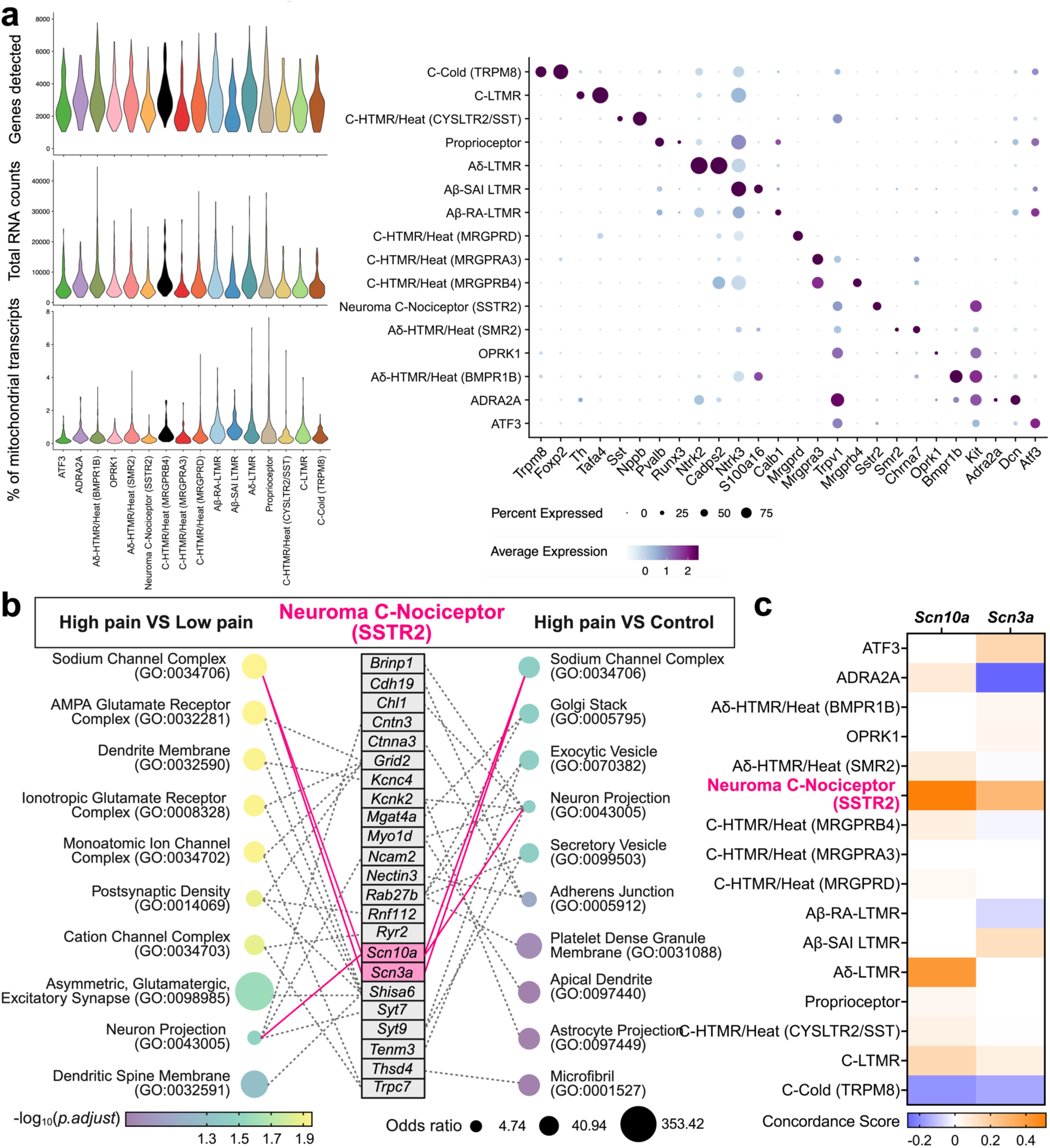
| Transcriptomic profiling of DRG across pain conditions. **a**, Violin plots showing quality control metrics, including the number of genes detected per nucleus (nFeature cutoff = 1,000), total RNA counts, and percentage of mitochondrial transcripts, and dot plots showing expression of canonical marker genes across neuronal populations. Dot size represents the percentage of nuclei expressing the gene, and dot colour reflects average expression level. Cell-type annotations follow previously described criteria^30^. **b**, Gene Ontology enrichment within the Neuroma C-Nociceptor (SSTR2) population. Left: top 10 Cellular Component terms for high-pain vs. low-pain; right: top 10 Cellular Component terms for high-pain vs. control, ranked by adjusted *P* values. Middle column lists DEGs contributing to each term; solid magenta lines connect genes shared between contrasts, gray dashed lines connect genes appearing in only one contrast. **c**, Concordance scores for *Scn10a* and *Scn3a* across sensory neuron subtypes. The score is positive when a gene is up-regulated in both comparisons (high-pain vs. low-pain and high-pain vs. control), negative when down-regulated in both, and zero when regulation is inconsistent or observed in only one comparison; larger magnitude indicates stronger concordance (see Methods for formula).

**Extended Data Fig. 7.**
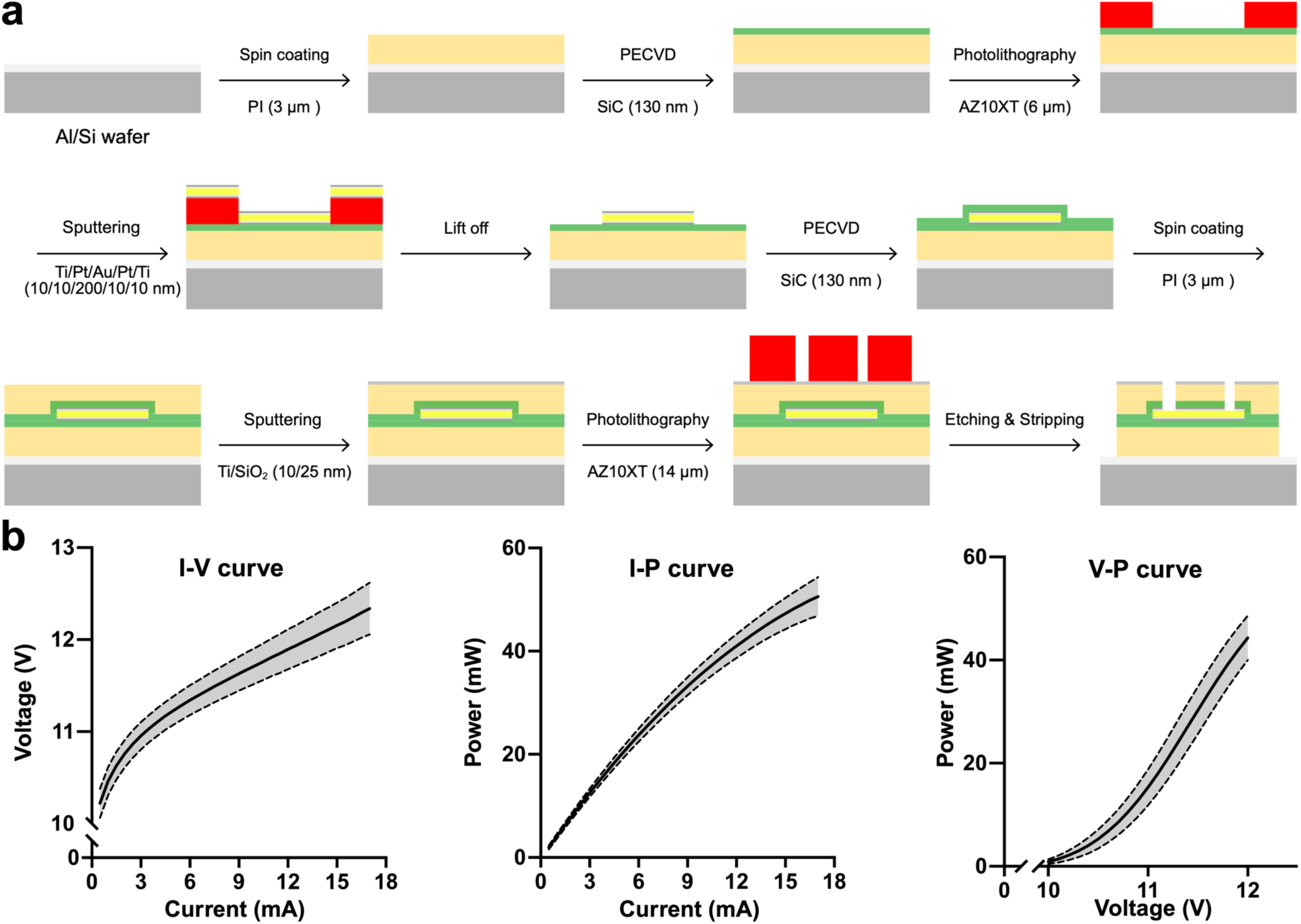
| Fabrication and characterization of the miniaturized LED cuff. **a**, Schematic of the fabrication process. A 3-µm polyimide (PI) layer is spin-coated onto an aluminum (Al)/silicon (Si) wafer, serving as a sacrificial substrate, and baked to complete imidization. A 130-nm silicon carbide (SiC) layer is deposited by plasma-enhanced chemical vapor deposition (PECVD). Metal interconnects consisting of Titanium/Platinum/Gold/Platinum/Titanium (Ti/Pt/Au/Pt/Ti) (10/10/200/10/10 nm) are sputtered and patterned by lift-off, followed by deposition of a second SiC layer and an additional PI layer. A Ti/SiO_2_ bilayer (10/25 nm) is sputtered on the surface, and final device patterning is performed by inductively coupled plasma–reactive ion etching (ICP-RIE) to define cuff geometry and electrode windows. **b**, Electrical and optical performance of the integrated blue micro-LEDs (470 nm). Representative current–voltage (I–V), current–power (I–P), and voltage–power (V–P) curves. Solid lines and shaded bands represent mean ± SEM across n = 24 devices.

**Extended Data Fig. 8.**
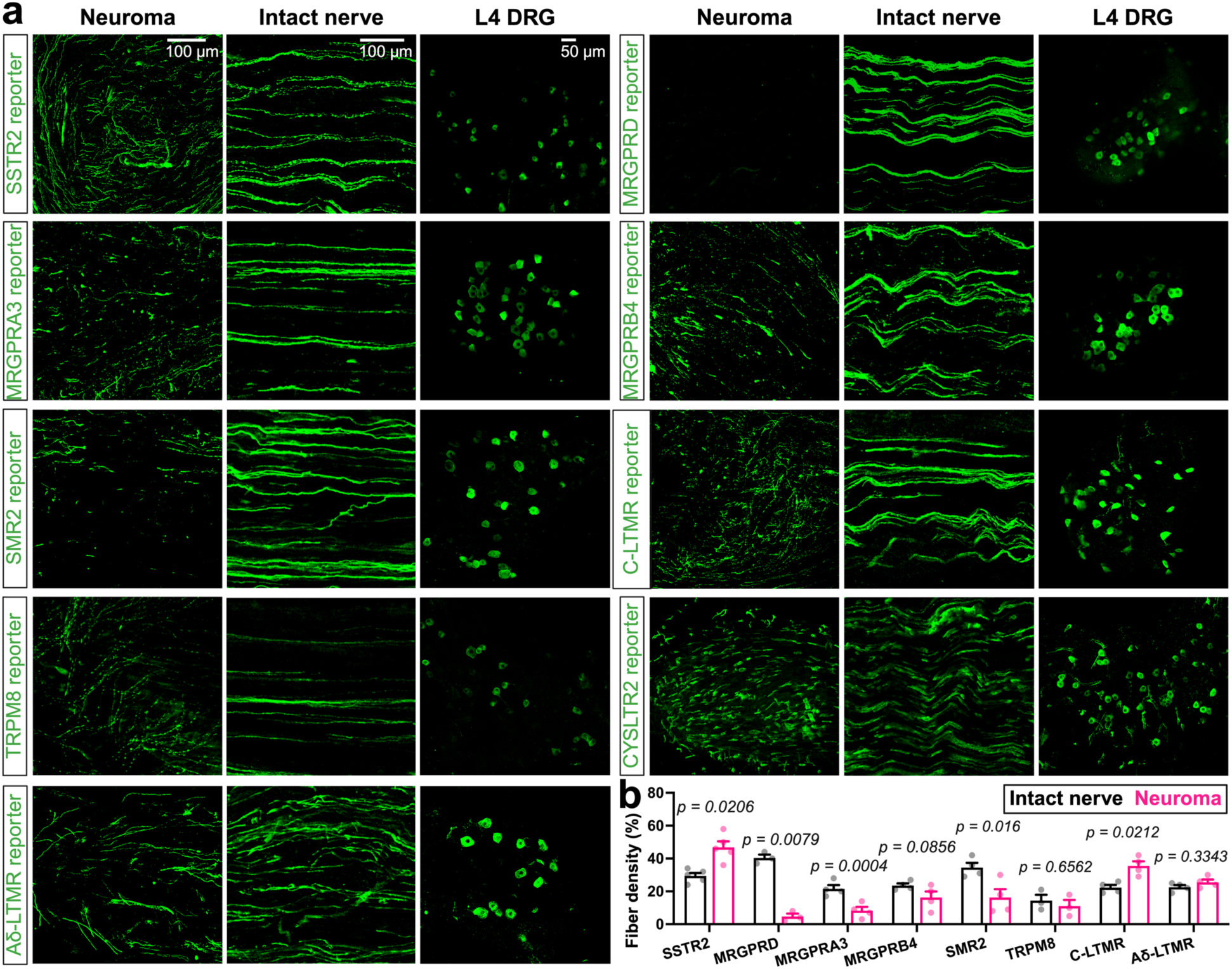
| Differential density of sensory fibre subtypes in neuromas. **a**, Representative images of reporter expression (ReaChR::mCitrine) in neuromas, contralateral intact nerves, and L4 DRG neurons from transgenic lines, including Neuroma C-Nociceptors (SSTR2), C-HTMR/Heat (MRGPRD), C-HTMR/Heat (MRGPRA3), C-HTMR/Heat (MRGPRB4), Aδ-HTMR/Heat (SMR2), C-Cold (TRPM8), C-LTMR, and Aδ-LTMRs. **b**, Quantification of fibre density, expressed as the percentage of axon-occupied area. Note that the increased density in neuromas labelled by the C-LTMR reporter may reflect contributions from both C-LTMRs and Th⁺ sympathetic nerves. C-HTMR/Heat (CYSLTR2/SST) fibres were not quantified because the transgenic line (*Cysltr2^Cre^*) labels a mixture of axons and non-neuronal cells in peripheral nerves. Data are mean ± SEM; each dot represents one nerve/neuroma. *P* values from two-sided paired t-tests are shown for each population. All these histology studies were performed following the optogenetic activation experiments at 8–12 weeks post-SNT.

**Extended Data Fig. 9.**
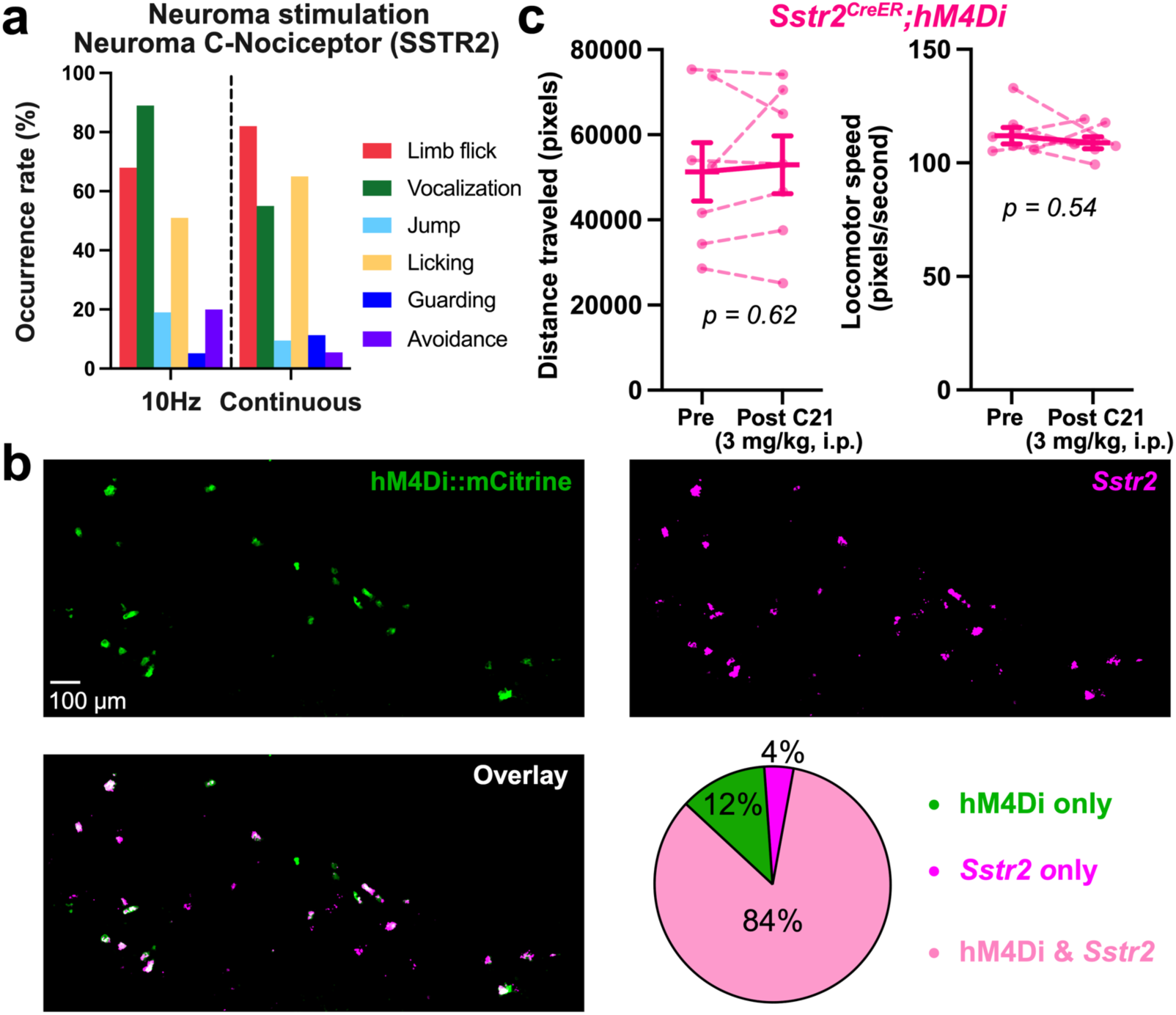
| Gain- and loss-of-function analyses of Neuroma C-Nociceptors (SSTR2). **a**, Occurrence of behaviours in response to a 5-s stimulation, delivered either as a 10 Hz train of 10-ms pulses or as continuous light. **b**, Representative images and quantification of combined *in situ* hybridization for *Sstr2* and immunolabeling of hM4Di::mCitrine in the ipsilateral L4 DRG of *Sstr2^T2a-CreER^; R26^LSL-hM4Di::mCitrine^* mice. Twenty L4 DRG sections from two *Sstr2^T2a-CreER^; R26^LSL-hM4Di::mCitrine^*mice were used to quantify the overlap ratio between *Sstr2* mRNA and hM4Di::mCitrine expression. **c**, Quantification of distance traveled and locomotor speed before and after acute chemogenetic silencing using a previously reported behavioural platform^53^. Data are mean ± SEM; each dot represents one mouse (n = 7 mice). *P* values from two-sided paired t-tests.

